# Anti-psychotic therapeutics modify gut microbial metabolism and modulate susceptibility to gastrointestinal infection in mice

**DOI:** 10.64898/2026.09.15.745804

**Authors:** C. Anthony Gacasan, Jaclyn Weinberg, Lyndsey D. Lipson, Gabrielle Webster, Dean P. Jones, Timothy R. Sampson, Rheinallt M. Jones, Michael H. Woodworth

**Affiliations:** Division of Gastroenterology, Hepatology and Nutrition Department of Pediatrics, Emory University School of Medicine, Atlanta, GA, USA; Division of Pulmonary, Allergy, Critical Care and Sleep Medicine, Department of Medicine, Emory University School of Medicine, Atlanta, GA; Department of Cell Biology, Emory University School of Medicine, Atlanta, GA 30322, United States; Division of Infectious Disease, Department of Medicine, Emory University School of Medicine, Atlanta, GA, USA

**Keywords:** Anti-Psychotics, Microbiome

## Abstract

Antipsychotic medications are widely prescribed and associated with increased infection risk, but underlying mechanisms remain unclear. We examined if antipsychotic-induced microbiome and metabolome alterations impair colonization resistance. Using a murine model, we evaluated haloperidol, olanzapine, risperidone, and quetiapine through longitudinal microbiome profiling, untargeted metabolomic fingerprinting, behavioral testing, and enteric infection challenge. Antipsychotic exposure induced persistent behavioral changes and increased susceptibility to *Citrobacter rodentium*, with quetiapine and olanzapine producing the greatest weight loss. Microbiome sequencing revealed treatment-specific shifts in beta diversity without consistent alpha diversity changes, while untargeted metabolomic analysis demonstrated robust, drug-specific metabolic reprogramming, particularly in lipid and sterol pathways. Although microbial compositional changes did not fully account for functional outcomes, their integration with metabolomics data revealed disrupted bacterial taxa-metabolite networks and loss of homeostatic metabolic modules. These results provide a potential mechanistic link between the disruptive effects of antipsychotics on the microbiome and impaired colonization resistance.

## Introduction

Antipsychotic medications are widely prescribed across psychiatric and non-psychiatric indications, including schizophrenia, bipolar disorder, delirium, and behavioral symptoms of dementia, with more than 23 million individuals affected by schizophrenia alone globally ^1^. Use of these agents is particularly common in older adults and long-term care facilities (LTCFs), where antipsychotics are frequently prescribed off-label and often chronically ^2, 3^. Across diverse populations and drug classes, antipsychotic use has been consistently associated with increased risk of infectious outcomes, including pneumonia, infection-related hospitalization, and mortality ^4–6^. Despite the robustness of these epidemiologic associations, the mechanisms underlying antipsychotic-associated infection susceptibility remain poorly defined, limiting the development of effective prevention strategies for patients committed to long-term therapy.

The intestinal microbiota plays a central role in host defense against enteric pathogens through colonization resistance. Furthermore, specific microbial metabolic functions that contribute to the strength of colonization resistance include generation of bacterially derived metabolites and the conversion and biotransformation of primary and conjugated bile acids (BAs) into secondary and deconjugated species, as well as lesser-studied metabolites such as bacterial bile acid amidates ^7–11^. These functions are also recognized as modulators of gut inflammation and risk of infection ^7, 8, 12^. While anaerobic antibiotics are among the most potent microbiome-disrupting exposures, many non-antibiotic medications have increasingly recognized effects on gut microbial communities, including antipsychotics ^13, 14^.

Antipsychotic medications (e.g., haloperidol, olanzapine) have been shown to inhibit bacterial growth in vitro, reduce alpha diversity in complex microbial communities, and increase bacterial resistance to antibiotics through induction of efflux pump expression ^12–14^. If the established epidemiologic association between antipsychotic use and infection risk is mediated, in part, through microbiota disruption, this could open new microbiota-directed prevention strategies for millions of patients receiving antipsychotics ^15, 16^. However, the in vivo microbiome-disrupting effects of clinically relevant antipsychotics and their functional consequences for infection susceptibility remain unclear.

In this study, we address these gaps through in vivo exposure of mice to haloperidol, olanzapine, risperidone, and quetiapine. We hypothesized that antipsychotic medications disrupt gut microbiota composition in vivo, modulate specific microbial metabolic functions, and ultimately contribute to increased susceptibility to pathogen colonization and infection. To that end, we investigated antipsychotic-induced effects on the gut microbiota through longitudinal analysis of fecal samples, assessed functional consequences using open-field behavioral testing following drug cessation, and evaluated colonization susceptibility using a *Citrobacter rodentium* model of enteropathogenic *Escherichia coli* ^17, 18^. To provide mechanistic insight, we employed both targeted and untargeted metabolomics approaches ^19, 20^. By integrating longitudinal microbial sequencing, metabolomics, and pathogen challenge experiments, this work elucidates an understudied consequence of antipsychotic exposure and supports a growing paradigm in which non-antibiotic medications exert microecological effects with meaningful consequences for infection risk and patient outcomes ^13^.

## Results

### Three weeks of antipsychotic use is sufficient to drive lasting behavioral changes even after therapeutic cessation

To investigate the effects of antipsychotic use on mice, we employed a one-month study in which specific pathogen-free (SPF) C57BL/6 mice were subjected to daily treatment with one of four antipsychotic therapeutics, haloperidol, quetiapine, risperidone, and olanzapine, or a vehicle control, using dosing paradigms consistent with previously described murine studies ^21–24^. Following 26 days of treatment, mice were transferred to our murine behavioral testing facilities and allowed a 4-day accommodation period and antipsychotic washout before undergoing open-field behavioral testing **(Figure 1A).** Although not reaching the statistical significance threshold, all antipsychotic treatment groups demonstrated shorter time spent in the center of the field after 10 minutes of observation. **(Figure 1B and F)** After 30 minutes of observation, most groups normalized closer to the vehicle control, with only the quetiapine treatment group maintaining relatively lower center time compared to vehicle, although this difference remained non-significant. **(Figure 1C and F)** Similarly, with the exception of the risperidone treatment group, all antipsychotic treatments resulted in decreased total distance traveled relative to the vehicle control group after 10 minutes of observation, with significant differences observed in the haloperidol group (p = 0.0342) and the olanzapine treatment group (p = 0.0306, **Figure 1D and F**). After 30 minutes of observation, most groups again maintained a shorter average total distance traveled compared to vehicle control, especially mice that had received olanzapine treatment. **(Figure 1E and F).** These relationships were preserved when center entries were analyzed as counts and when normalized to distance traveled, a proxy for changes in murine “anxiety” behavior **(Supplementary Figure 1).**

**Figure 1.**
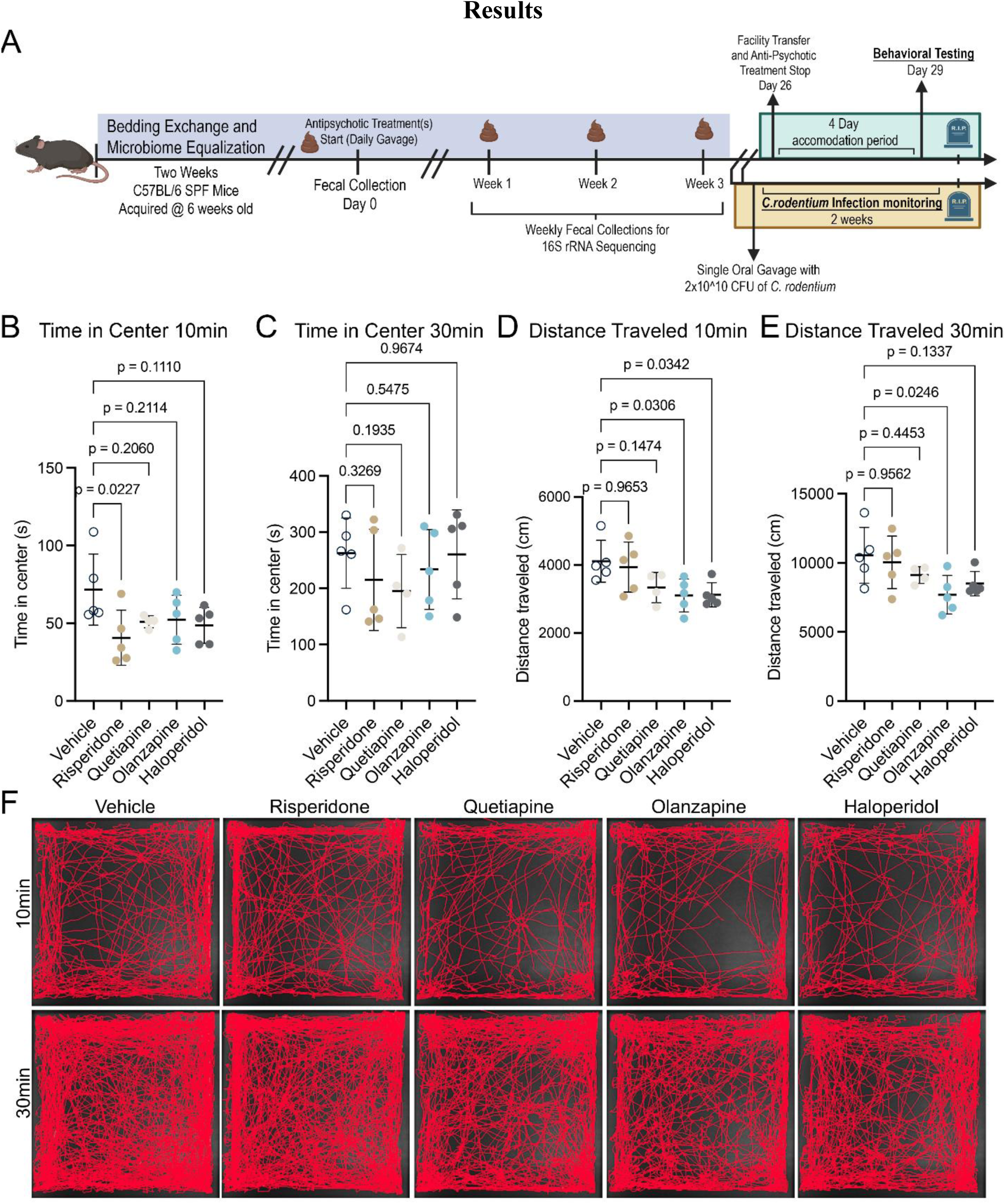
Antipsychotic treatment leads to lasting behavioral changes after therapeutic cessation. **(A)\*** Schematic of Experimental Design, n=5 per treatment group **(B)** Scatter dot plot showing time spent in the center of the open field after 10 minutes of observation, presented as group means with standard deviation, post-hoc test is Dunnett’s multiple comparison test. **(C)** Time spent in the center of the open field after 30 minutes of observation. **(D)** Scatter dot plot of total distance traveled during the first 10 minutes of open field testing. Statistical significance reflects pairwise comparisons derived from ANOVA. **(E)** Total distance traveled after 30 minutes of observation. **(F)** Representative open field trajectories for each antipsychotic treatment group. [*Created with BioRender.com (Gacasan, A., 2026; https://BioRender.com/0imn4ww) and subsequently modified.]

### Antipsychotic therapeutic administration promotes gastrointestinal colonization of pathogenic *C. rodentium*

To determine the functional changes to the gut and as a proxy to gastrointestinal infection and colonization risk due to antipsychotic administration, we employed a *C. rodentium* model of gut infection, a common model used for enteropathogenic hemorrhagic *E. coli* infection ^17, 18^. After colonization with *C. rodentium*, all treatment groups exhibited infection-associated weight loss by two weeks post-infection, with vehicle-treated mice showing the mildest decline. Neither haloperidol nor risperidone significantly altered the rate of weight loss when compared to vehicle control (Haloperidol: Δslope = –0.0099 ± 0.0179, p-value = 0.91; Risperidone: Δslope = –0.0071 ± 0.0179, p-value = 0.96, **Figure 2A**). In contrast, both Quetiapine and Olanzapine produced significantly steeper rates of weight loss (Quetiapine: Δslope = –0.0507 ± 0.0179, p-value = 0.018; Olanzapine Δslope = –0.0501 ± 0.0179, p-value = 0.020). Additionally, to assess relative bacterial load, colony forming units (CFU)/gram of stool was assessed via selective culture. (**Figure 2B and C)**. Linear ANCOVA revealed only Risperidone demonstrated statistical significance when compared to vehicle control (Δslope = 0.263 ± 0.0695, P-adj = 0.00141). While the rate of CFUs/g of stool was not significant for Olanzapine when compared to vehicle control (Δslope = 0.142 ± 0.0719, P-adj = 0.157) it did demonstrate a significant difference to vehicle at Day 14 Post-Infection (P-adj = 0.0468). These findings, together with persistent behavioral changes following a 4-day washout, suggest lasting physiological alterations not solely attributable to the direct effects of antipsychotic exposure, potentially implicating the gut microbiome. (**Supplementary Table 1**).

**Figure 2.**
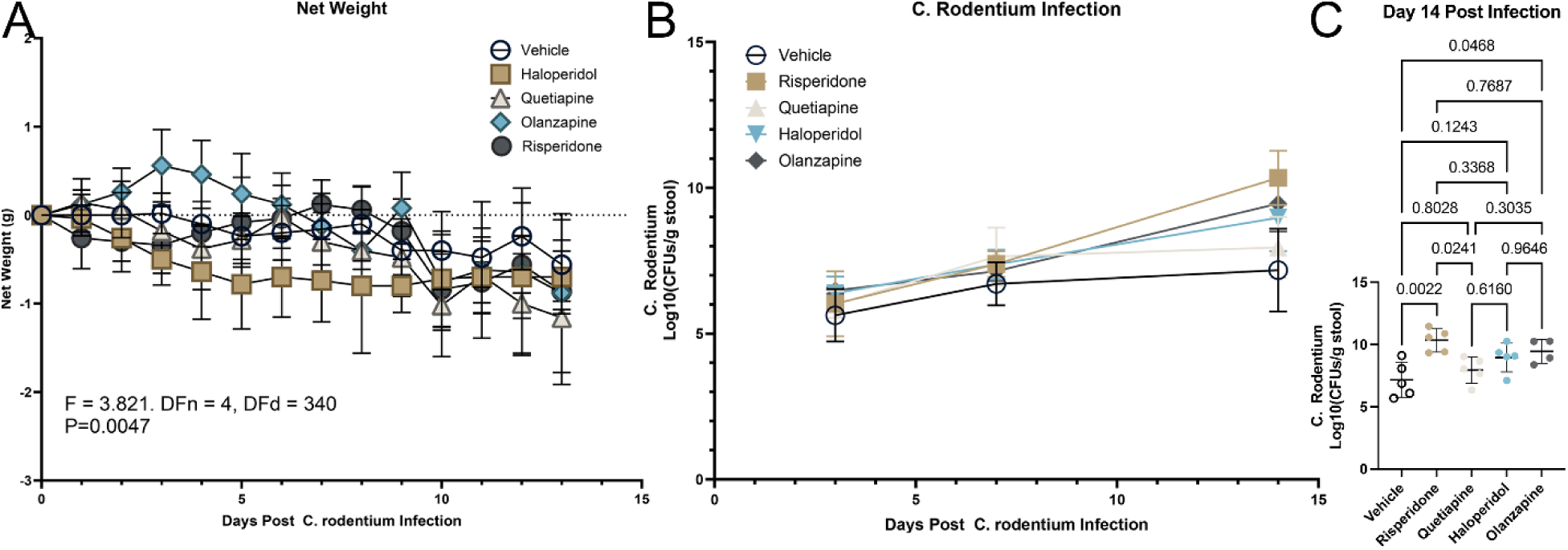
Antipsychotic treatment alters host weight trajectories and increases *C. rodentium* burden following oral inoculation. (**A)** Net weight change over the course of infection, measured from day 0 (first day of gavage) through necropsy at two weeks post–*C. rodentium* inoculation. Data are shown as mean ± SEM, with repeated-measures ANOVA statistics indicated**. (B)** Fecal *C. rodentium* burden over time, quantified as log₁₀ colony-forming units (CFU) per gram of stool across the two-week infection period. (**C)** *C. rodentium* burden at day 14 post-infection, shown as dot plots with mean ± SD. Statistical comparisons between treatment groups were performed using ANOVA with pairwise post hoc testing using Dunnet’s; exact p-values are shown.

### Antipsychotic administration modifies gut microbial diversity

Microbial community processing and diversity analyses were performed using established QIIME-based workflows and phylogenetic methods of non-challenged antipsychotic treated mice to assess the effects of antipsychotics on microbiome composition ^25, 26^. Beta-diversity analysis demonstrated that the temporal trajectory of gut microbial community structure differed significantly among treatment groups. PERMANOVA revealed significant treatment group-by-timepoint interactions for both Bray–Curtis dissimilarity (Df = 12, R² = 0.164, F = 1.91, p < 0.001) and Jaccard distance (Df = 12, R² = 0.139, F = 1.22, p = 0.001), indicating that antipsychotic exposure was associated with treatment-specific changes in microbial community composition over time (**Figure 3A**). To summarize longitudinal trends in alpha diversity across the treatment period, linear slopes were estimated within each group. Observed richness declined significantly over time within the olanzapine group (slope = –678.3, p < 0.05; **Figure 3B**), whereas no significant temporal trends were detected in the other treatment groups. No significant temporal changes were detected in Shannon or Simpson diversity in any group (**Figure 3C and D**). Collectively, these analyses indicate that changes in microbial community composition over time varied among treatment groups, while a significant decline in richness was detected only within the olanzapine-treated group.

**Figure 3.**
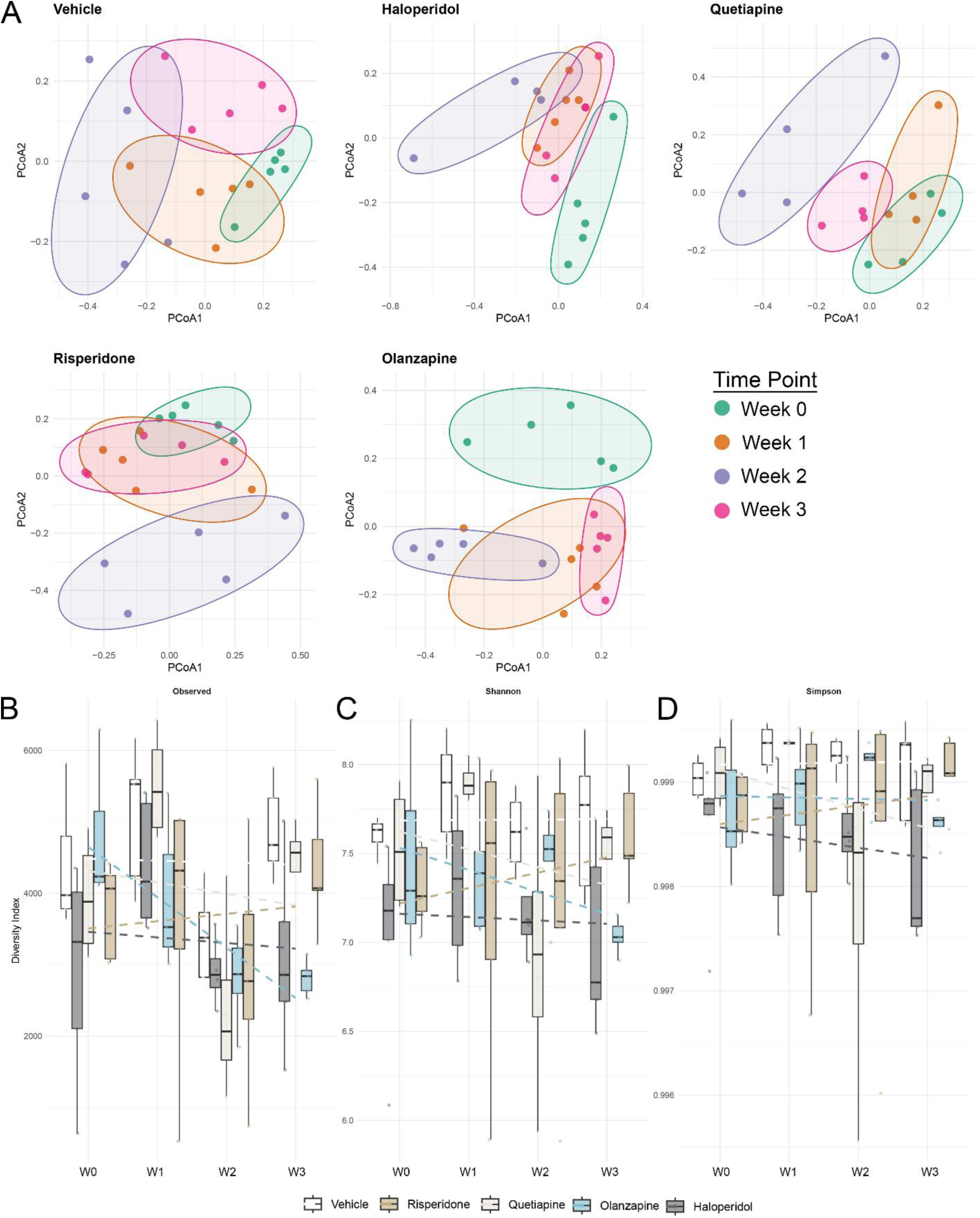
Antipsychotic treatment drives a variable fecal microbiome composition over time. **(A)** Principal coordinates analysis (PCoA) of gut microbial community structure stratified by treatment group. Bray–Curtis dissimilarities were calculated from rarefied 16S rRNA gene sequencing data and visualized using PCoA for each treatment group independently. Each panel represents a single treatment group (Vehicle, Haloperidol, Quetiapine, Risperidone, or Olanzapine), with points colored by sampling time point (Weeks 0-3).**(B-D)** Alpha diversity dynamics across antipsychotic treatment groups over time. **(B)** Observed ASV richness, **(C)** Shannon diversity, and **(D)** Simpson diversity indices were calculated from 16S rRNA gene sequencing data and plotted for each treatment group at each time point. Boxplots represent the distribution of diversity values within each treatment group at each time point, with individual points indicating single samples. Dashed lines depict linear trend fits across time for each treatment group. Facets are shown separately for each diversity metric with free y-axis scaling.

### Differential abundance analysis reveals treatment-specific microbiome compositional changes

Across all treatment groups and time points, microbial communities were dominated by *Bacillota* (formerly *Firmicutes*). At the phylum level, risperidone was associated with a transient reduction in *Bacillota* and corresponding increases in *Bacteroidata* and *Pseudomonadota*, while *Verrucomicrobiota* transiently increased with quetiapine and risperidone but remained comparatively low and stable with vehicle and olanzapine treatment (**Figure 4A**). At the genus level, the microbiota was dominated by *Lactobacillus*, *Faecalibaculum*, *Dubosiella*, *Lachnospiraceae* NK4A136 group, *Lachnoclostridium*, and several Eubacterium groups (Figure 4B). Vehicle-treated mice maintained relatively high *Dubosiella* abundance and exhibited only modest changes in Lactobacillus, whereas all antipsychotic-treated groups demonstrated progressive Lactobacillus expansion, reaching approximately 33–48% with haloperidol, 25–31% with olanzapine, 31–42% with quetiapine, and 16–30% with risperidone at later time points. *Akkermansia* remained relatively low across groups but showed transient increases with quetiapine and risperidone, consistent with prior reports linking antipsychotic exposure to enrichment of lactic acid bacteria and shifts in mucin-associated taxa (**Figure 4B**) ^12, 14, 24^.

**Figure 4.**
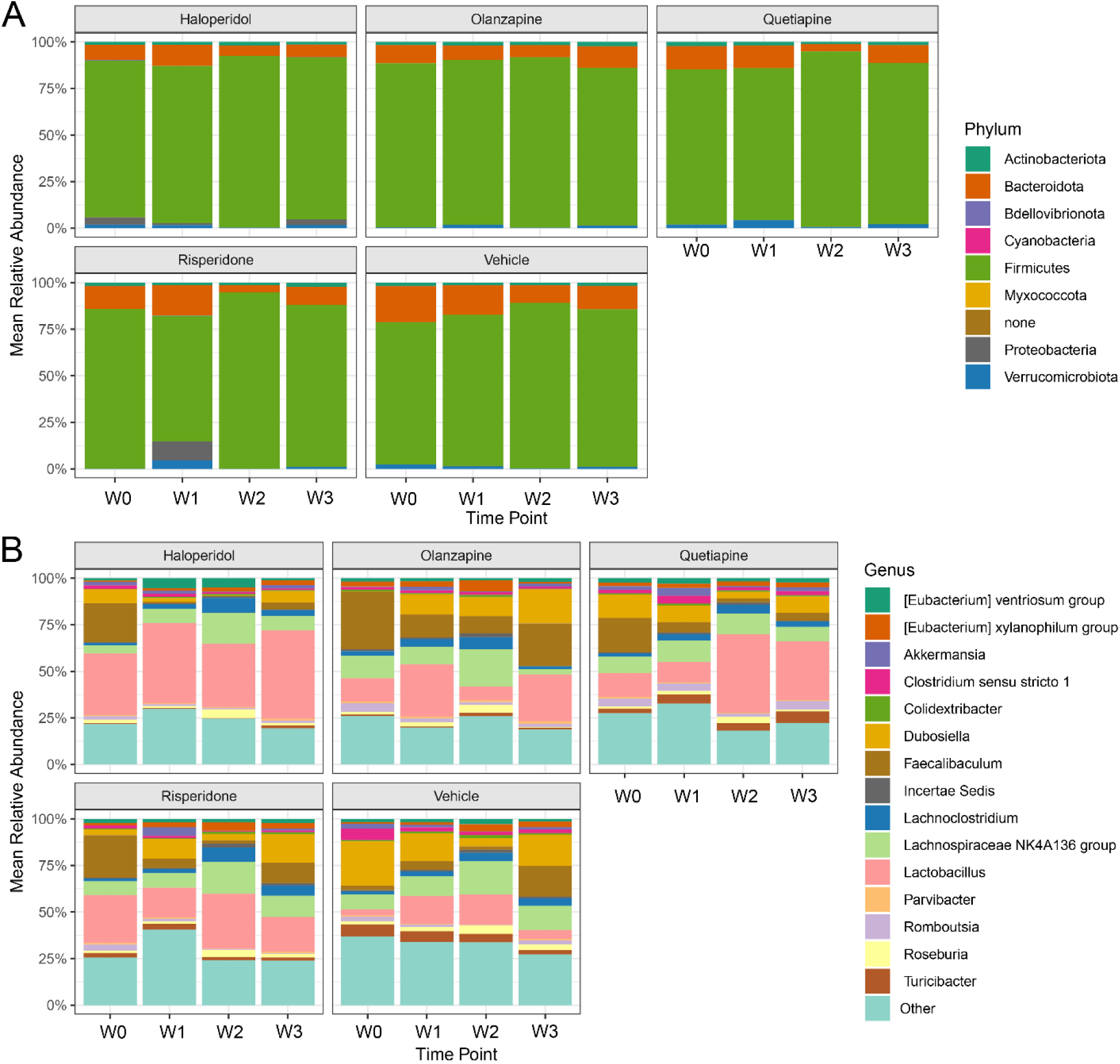
Antipsychotic treatment is associated with treatment-specific and time-dependent alterations in gut microbial composition. **(A)** Phylum-level and **(B)** genus-level mean relative-abundance profiles for each treatment group across time points. For each mouse, taxonomic abundances were normalized to the total number of classified sequences such that the relative abundances summed to 100%. Relative abundances were then averaged across mice within each treatment group and time point. Each stacked bar therefore represents the group mean compositional profile and sums to 100%

Although alpha-diversity measures were largely stable across treatment groups, longitudinal analyses demonstrated treatment-specific changes in several dominant genera, indicating that antipsychotic exposure altered community composition without uniformly changing within-sample diversity. Temporal trajectories of the five dominant genera were evaluated using longitudinal generalized estimating equations with mouse as the repeated-measures cluster and arcsine-square-root-transformed relative abundances. Benjamini–Hochberg FDR-adjusted pairwise whole-trajectory Wald tests showed that Faecalibaculum, *Dubosiella*, and *Lactobacillus* trajectories differed between Vehicle and each antipsychotic treatment [Faecalibaculum: Wald χ²(3) = 53.7–208.6, all q ≤ 1.83 × 10⁻⁸; Dubosiella: Wald χ²(3) = 10.4–105.5, all q ≤ 0.023; Lactobacillus: Wald χ²(3) = 13.2–117.0, all q ≤ 0.0096]. *Dubosiella* exhibited the greatest treatment specificity, with significant differences among all drug pairs except quetiapine and haloperidol. Quetiapine also produced a distinct Lactobacillus trajectory relative to risperidone, olanzapine, and haloperidol, whereas the haloperidol-associated *Faecalibaculum* trajectory differed from those observed with risperidone and quetiapine. *Lachnoclostridium* showed a more selective response, with olanzapine differing from both Vehicle and risperidone, while no antipsychotic significantly altered the *Lachnospiraceae* NK4A136 group trajectory relative to Vehicle after FDR correction (**Supplementary Figure 2, Supplementary Table 2**).

### Antipsychotic treatment drives a distinct metabolomic signature

To assess functional consequences of antipsychotic exposure, we performed untargeted high-resolution metabolomics analysis of fecal pellets collected before and after the three weeks of antipsychotic exposure, using established LC-MS feature detection and putative annotation workflows ^19, 20, 27^. Antipsychotic administration resulted in a distinct fecal metabolome across treatment groups, with all groups exhibiting discrete separation by PLS-DA and pairwise PERMANOVA revealing strong, drug-specific differences between treatments; all pairwise comparisons excluding haloperidol versus risperidone demonstrated FDR-adjusted significance (P-adj < 0.05, **Figure 5A, Supplementary Table 3**). Following integration of high-confidence (score = 3) putative annotations from xMSannotator and unsupervised clustering, olanzapine exhibited metabolomic profiles most similar to vehicle control, whereas risperidone and haloperidol clustered together, and Quetiapine formed a distinct cluster (**Figure 5B**).

**Figure 5.**
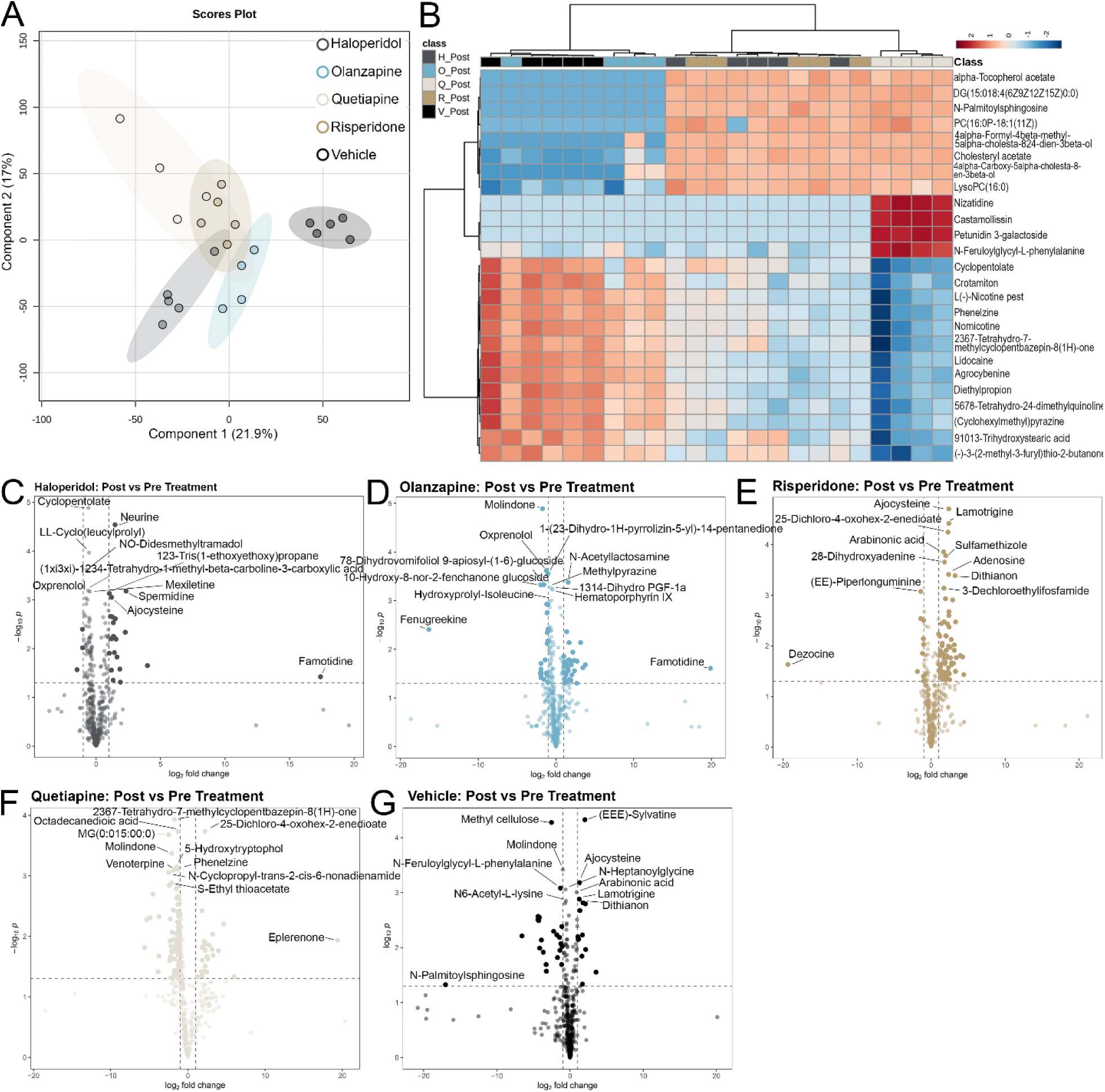
Antipsychotic treatment induces distinct metabolomic signatures prior to *C. rodentium* infection. **(A)** Unsupervised principal component analysis (PCA) of untargeted fecal metabolomics profiles collected after three weeks of antipsychotic treatment and prior to *C. rodentium* infection. Samples are colored by treatment group, illustrating treatment-associated separation in metabolic composition**. (B)** Heatmap of the top 25 putatively annotated metabolites identified using xMSannotator, with samples clustered by treatment class. Metabolites and samples were hierarchically clustered using Euclidean distance and Ward’s linkage method. **(C–G)** Volcano plots depicting post-treatment versus pre-treatment differential metabolite abundance for each group: **(C)** Haloperidol, **(D)** Olanzapine, **(E)** Risperidone, **(F)** Quetiapine, and **(G)** Vehicle. Points represent putatively annotated metabolic features plotted as log₂ fold change (Post/Pre) versus −log₁₀(p-value), with selected metabolites being those most differentially enriched based on p-value and fold change. All annotations remain putative and may represent isomers of the generated annotation.

One-way ANOVA identified several metabolites significantly differentiated across antipsychotic treatment groups following FDR correction. These features spanned multiple chemical classes, including xenobiotic-associated compounds, lipid species, microbial- and diet-derived metabolites, and neuroactive metabolic intermediates (**Figures 5C-G**). Among the putatively annotated features were xenobiotic-related metabolites such as nicotine-associated alkaloids (nornicotine, arecoline) and monoamine-related molecules including kynuramine, metanephrine, and tetrahydroharmol. Although mice were only administered antipsychotics without any additional concomitant medications, several putatively annotated xenobiotic-like compounds (e.g., lidocaine, mexiletine, tocainide) also showed significant differences across treatment groups. Significant alterations were observed in lipid mediators and membrane-associated species, including prostaglandin E2 (PGE2), 19-hydroxy-PGE2, diacylglycerols, phosphatidylcholines, lysophosphatidylcholines, and cholesterol esters. In addition, several flavonoid glycosides and phenolic conjugates exhibited significant group-wise differences. Metabolites within the tryptophan–kynurenine and indole-derived pathways, including kynuramine and tetrahydroharmol, were also significantly altered across treatments, consistent with the established role of microbial amino acid metabolism in shaping host metabolic and immune tone (**Supplementary Table 4**) ^9, 10^.

### Pathway Enrichment and Network based metabolomics reveals convergent metabolic modules

To delineate treatment-specific metabolic alterations, pathway enrichment analysis was performed using pairwise comparisons between vehicle and each antipsychotic treatment on untargeted fecal metabolomics data. Enrichment analysis using mummichog identified both shared and treatment-specific metabolic pathways relative to vehicle ^19, 20, 27^ (**Figure 6A-D)**. Across all antipsychotic treatment groups, lipid- and sterol-associated pathways were consistently enriched, with squalene and cholesterol biosynthesis pathways enriched across risperidone, quetiapine, and haloperidol treatment groups. Additional pathways commonly enriched included glycerophospholipid metabolism, fatty acid–associated pathways, vitamin metabolism, and cytochrome P450– associated drug metabolism. Pathways related to prostaglandin formation from arachidonate and glycosphingolipid metabolism were also enriched across multiple treatment groups, indicating convergent alterations in lipid-associated metabolic networks within the gut.

**Figure 6.**
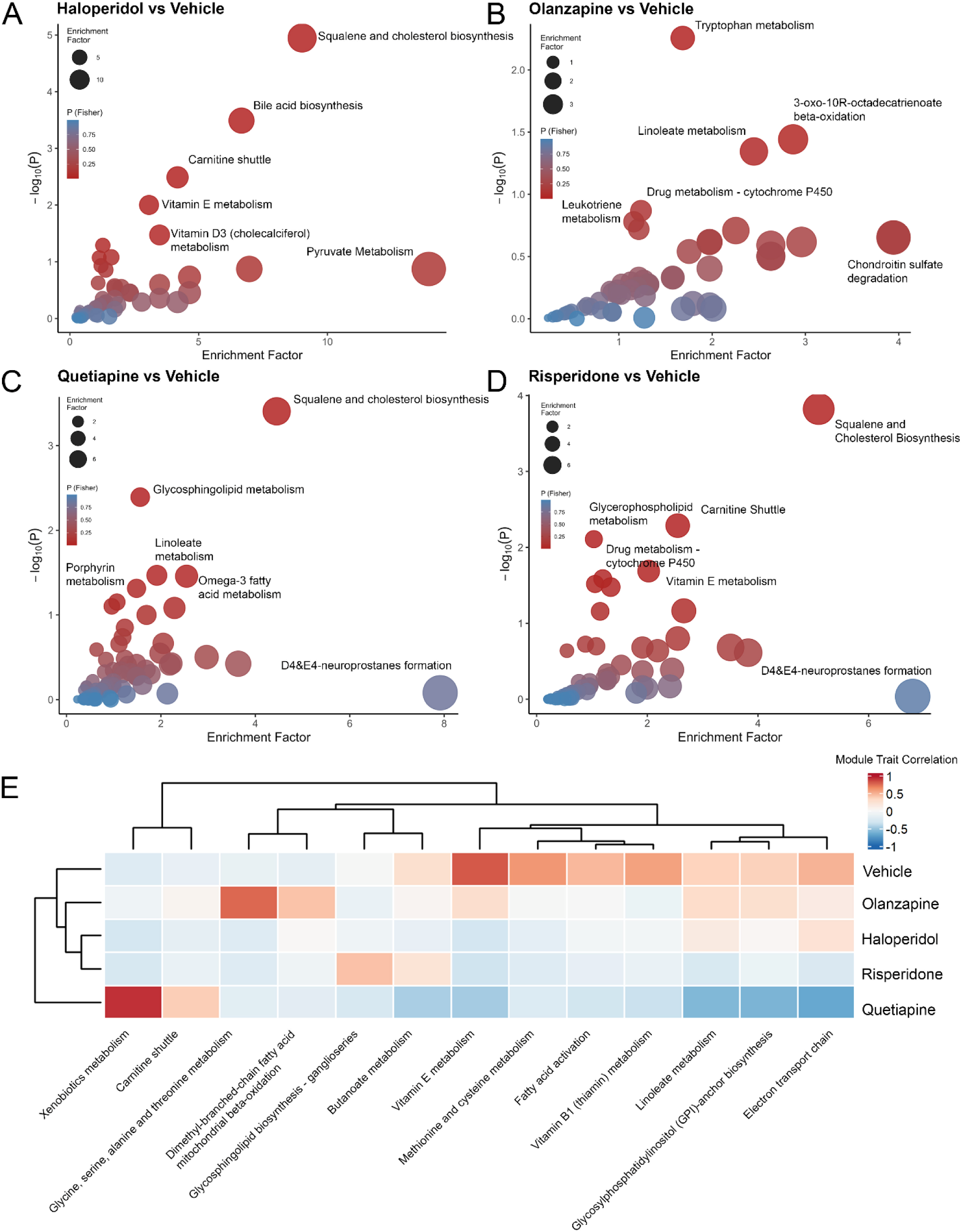
Antipsychotic treatment induces shared and drug-specific metabolic pathway perturbations revealed by pathway enrichment and network-based analyses. **(A-D)** Mummichog pathway enrichment analysis of untargeted metabolomics data comparing each antipsychotic treatment to vehicle. Bubble plots display enriched metabolic pathways for **(A)** haloperidol, **(B)** olanzapine, **(C)** quetiapine, and **(D)** risperidone relative to vehicle. The x-axis represents the enrichment factor, the y-axis denotes −log₁₀(p-value), and bubble size reflects enrichment factor. Bubble color indicates pathway significance, with red representing more significant enrichment and blue representing less significant enrichment. All analyses were performed using pairwise mummichog comparisons between each antipsychotic and vehicle control. **(E)** Weighted correlation network analysis (WGCNA) of untargeted metabolomics data using a minimum module size of 250 features. Heatmap of module–trait correlations between metabolite modules and treatment groups. Each treatment group was modeled as a separate binary trait, with samples in that group coded as 1 and all other samples coded as 0; therefore, correlations represent each treatment group relative to all remaining groups combined, and no single baseline or reference group was used. Color intensity reflects the strength and direction of the correlation (blue, negative; red, positive). Rows represent treatment groups, and columns represent metabolite modules, labeled by their most significantly enriched pathway identified by mummichog.

The consistent enrichment of lipid- and sterol-associated pathways across pharmacologically distinct antipsychotics suggests a shared metabolic response among the drugs tested, potentially reflecting common physicochemical properties, host xenobiotic responses, or microbiome adaptations to chronic drug exposure. Collectively, these findings implicate the gut as a potential site of antipsychotic-induced dyslipidemia and altered inflammatory tone, consistent with established clinical and mechanistic links between antipsychotics and metabolic dysfunction ^28–30^.

Each antipsychotic also exhibited distinct pathway enrichment profiles. Haloperidol treatment demonstrated pronounced enrichment of sterol- and bile-associated pathways, including squalene and cholesterol biosynthesis and bile-acid biosynthesis, aligning with prior evidence that antipsychotics can influence sterol biosynthetic pathways ^22^. Additional enriched pathways included the carnitine shuttle, vitamin E, vitamin D3, and vitamin B6 metabolism, as well as amino-acid–related pathways such as tryptophan and tyrosine metabolism (**Figure 6A**). In contrast, olanzapine treatment exhibited a divergent enrichment profile dominated by amino-acid and vitamin metabolism pathways. Tryptophan metabolism was the most prominently enriched pathway, accompanied by enrichment of linoleate metabolism, leukotriene metabolism, lipoate metabolism, vitamin B6 and vitamin D3 metabolism, β-oxidation–related pathways, glycosphingolipid biosynthesis, and cytochrome P450–associated drug metabolism (**Figure 6B**). Quetiapine treatment showed broader enrichment of polyunsaturated fatty-acid metabolism, including linoleate and omega-3 fatty-acid metabolism, in addition to glycosphingolipid metabolism, porphyrin metabolism, vitamin E metabolism, biopterin metabolism, and cytochrome P450–associated drug metabolism (**Figure 6C**). Risperidone treatment was characterized by enrichment of lipid transport and signaling pathways, including the carnitine shuttle, glycerophospholipid metabolism, and glycosphingolipid metabolism, along with enrichment of tyrosine metabolism and prostaglandin formation from arachidonate (**Figure 6D**). Together, these pathway analysis findings indicate that the four tested antipsychotics had distinct mechanistic effects that converged on disrupted lipid-sterol metabolism; risperidone through lipid signaling, quetiapine through polyunsaturated fatty-acid metabolism, haloperidol through squalene/cholesterol and bile acid signaling, and olanzapine through amino-acid and inflammatory signaling.

To identify coordinated metabolic programs associated with antipsychotic treatment, weighted gene correlation network analysis (WGCNA) was applied to the untargeted metabolomics data ^31, 32^. This network-based approach constructs modules of co-varying metabolites, enabling detection of higher-order metabolic structure not captured by univariate analyses. Integration of WGCNA with pathway enrichment analysis revealed several drug-specific module–trait associations. Olanzapine treatment showed strong correlations with modules enriched for glycine, serine, alanine, and threonine metabolism, as well as branched-chain β-oxidation pathways, suggesting a shift toward amino acid catabolism consistent with microbial metabolic remodeling described in prior literature ^9, 10^. Quetiapine treatment was distinct in that it exhibited strong negative correlations with most modules, indicating broad suppression of coordinated metabolic networks. The two modules uniquely positively correlated with quetiapine were enriched for xenobiotic metabolism and carnitine metabolism pathways (**Figure 6E)**. In contrast, the majority of metabolite modules were most strongly associated with vehicle-treated mice, reflecting coordinated metabolic programs maintained under physiologic conditions. These vehicle-associated modules were enriched for pathways central to gut metabolic homeostasis, including butanoate metabolism, fatty-acid activation, methionine and cysteine metabolism, linoleate metabolism, electron transport chain–associated metabolites, vitamin E and vitamin B1 metabolism, and GPI-anchor biosynthesis. The loss of these coordinated metabolic networks across antipsychotic treatment groups, consistent with pathway-level alterations identified in univariate analyses, suggests a shared disruption of the metabolic architecture of the gut dismantled by antipsychotics.

### Genus-resolved microbe–metabolite network architecture reveals structured associations

To link microbial community structure with gut chemical phenotypes, we constructed a bipartite association network between bacterial genera and annotated metabolites **(Figure 7A and B, Supplementary Figure 3)**^20, 31, 32^. This integrated analysis revealed that antipsychotic exposure was associated with coordinated alterations in gut microbial composition and fecal metabolic profiles, here we limited our analysis to a select few genera of interest.

**Figure 7.**
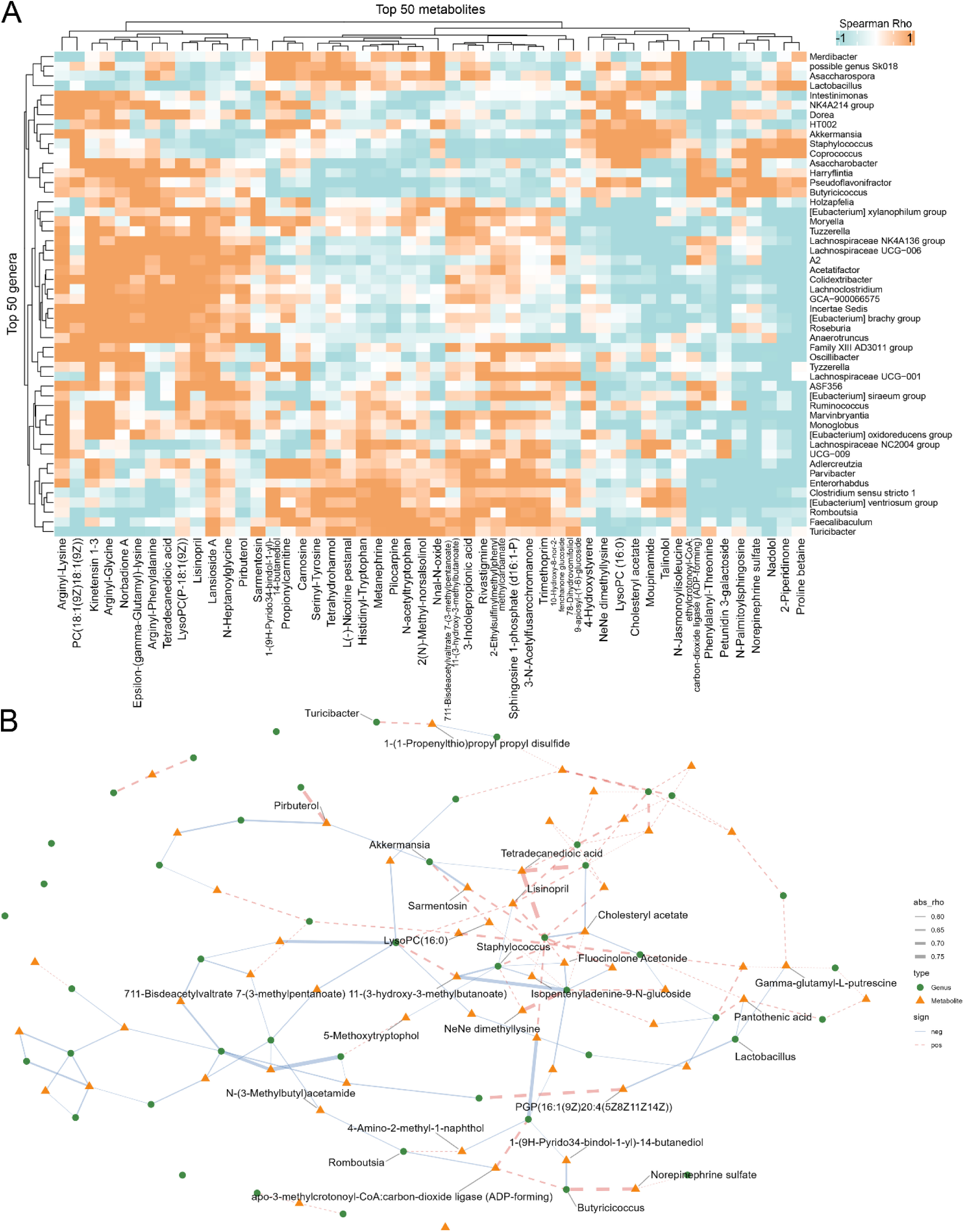
Integrated microbiome–metabolome associations after anti-psychotic treatment (pre–C. rodentium challenge). **(A)** Heatmap of Spearman correlation coefficients (ρ) between the top 50 bacterial genera (16S rRNA gene sequencing, CLR-transformed) and the top 50 putatively annotated untargeted metabolites (LC–MS) at the final antipsychotic treatment time point (week 3). Genera and metabolites were ranked by maximum absolute correlation magnitude across all pairwise associations. Colors represent Spearman’s ρ, with red indicating positive associations and blue indicating negative associations. **(B)** Bipartite genus–metabolite association network constructed from the strongest correlations used for visualization. Nodes represent genera (green circles) and metabolites (orange triangles). Edges are weighted by |ρ| (thicker = stronger association) and colored/typed by sign (positive = red dashed; negative = blue solid). Labels highlight the genera Akkermansia, Lactobacillus, Turicibacter, Butyriciococcus, Staphylococcus, Clostridium, and Romboutsia and their 1-hop putatively annotated metabolite neighbors.

Several *Lachnospiraceae*, which are commonly associated with gut health and short-chain fatty acid production, were correlated with homeostatic metabolites. For example, the *Lachnospiraceae NK4A136* group was positively associated with Norbadione A (ρ = 0.648) and ε-(γ-glutamyl)-lysine (ρ = 0.381), as well as lipid-associated metabolites including LysoPC(16:0) (ρ = 0.577) and (3a5b7a)-23-carboxy…glucuronide (ρ = 0.502). This genus also showed a modest positive correlation with sphingosine-1-phosphate (d16:1-P) (ρ = 0.140). These patterns are consistent with prior observations that *Lachnospiraceae* taxa contribute to gut metabolic homeostasis and short-chain fatty acid-associated metabolic networks ^10, 33, 34^.

Network analysis identified *Butyricicoccus* as a central hub within the neuro-metabolic axis, exhibiting some of the strongest correlations in the dataset (**Figure 7B, Supplementary Figure 3)**. Notably, *Butyricicoccus* was positively associated with norepinephrine sulfate (ρ = 0.734) and N-palmitoylsphingosine (ρ = 0.671), consistent with prior work linking butyrate-producing taxa to neurometabolic signaling ^33, 34^. Together with *Romboutsia*, which occupied a similar network neighborhood centered on apo-3-methylcrotonoyl-CoA:carbon-dioxide ligase, these taxa appear to define a coordinated microbial-metabolic module associated with host neuromodulatory tone. In contrast, *Butyricicoccus* was inversely associated with sphingosine-1-phosphate (d16:1-P) (ρ = −0.302) and 3-indolepropionic acid (ρ = −0.350), suggesting a shift in sphingolipid signaling and microbiota-derived neuroactive metabolites. Notably, sphingosine-1-phosphate is a bioactive lipid mediator implicated in cell survival, epithelial barrier integrity, and immune cell trafficking, raising the possibility that disruption of this microbial-metabolic axis may influence host protective signaling pathways^35, 36^. The importance of *Romboutsia* is further underscored by its selection in the supervised multivariate machine learning model DIABLO (block sPLS-DA), suggesting it may be a key driver of antipsychotic-associated metabolome remodeling (**Supplementary Figure 4**).

In contrast to these putatively homeostatic taxa, *Akkermansia* and *Staphylococcus* exhibited predominantly negative correlations across a broad range of annotated metabolites, particularly within lipid-associated pathways. Both taxa were inversely associated with sphingosine-1-phosphate (d16:1-P) (ρ = −0.185 and ρ = −0.484, respectively), while *Akkermansia* showed widespread negative correlations with lipid species and aromatic metabolites, including (3a5b7a)-23-Carboxy-7-hydroxy-24-norcholan-3-yl-b-D-Glucopyranosiduronic acid (ρ = −0.570) and 4-hydroxystyrene (ρ = −0.657) ^10^. Conversely, *Lactobacillus* and *Staphylococcus* were associated with enrichment of cholesteryl acetate (ρ = 0.349 and 0.642, respectively) and proline betaine (ρ = 0.449), while remaining negatively correlated with multiple lipid- and peptide-associated metabolites, including PGP(16:1/20:4) (ρ = −0.624) and arginyl-lysine (ρ = −0.535).

Taken together, these findings suggest that antipsychotic exposure not only shifts microbial composition, but also significantly reorganizes the structure of microbiome–metabolite interactions. Specifically, loss of butyrate-associated hub taxa such as *Butyricicoccus* is accompanied by a collapse of coordinated neurometabolic signaling networks and a concomitant emergence of taxa associated with broadly suppressive or dysregulated lipid-metabolic interactions. This network-level rewiring provides a possible biologic framework linking microbial perturbation to altered sterol, sphingolipid, and neuroactive metabolite signaling, which may underlie the impaired colonization resistance and behavioral phenotypes observed in antipsychotic-treated mice in addition to identifying candidate microbial-metabolic relationships that may be prioritized for future functional investigation.

## Discussion

In this study, we demonstrate that short-term exposure to four clinically relevant antipsychotics induces treatment-specific restructuring of the gut microbiome and fecal metabolome in mice, accompanied by increased suscdeptibility to *C. rodentium* colonization. These findings extend growing evidence that non-antibiotic medications can impair colonization resistance through microbiome effects^37^ and provide a link between antipsychotic use and the increased infection risk observed in epidemiologic studies ^3, 4, 6^.

Antipsychotic pretreatment increased *C. rodentium* colonization density across all four antipsychotics tested, with risperidone producing a significantly steeper trajectory compared to vehicle. Quetiapine and olanzapine additionally exacerbated infection-associated weight loss, suggesting impaired host resilience to enteric challenge. Open-field behavioral testing after a four-day washout revealed persistent reductions in locomotor activity, reaching significance for haloperidol and olanzapine at early time points, though most behavioral comparisons did not reach statistical significance. Because the washout period separates these phenotypes from the expected pharmacologic activity of the drugs studied, they are consistent with physiologic changes, potentially including microbiome-mediated effects, though the behavioral data alone cannot distinguish between central and peripheral mechanisms ^23^.

Longitudinal 16S rRNA gene amplicon sequencing revealed significant treatment-by-time interactions in beta diversity without uniform reductions in alpha diversity. Only olanzapine produced a significant decline in observed richness over time, while Shannon diversity measures remained stable across all treatment groups. This pattern of treatment-specific compositional restructuring, rather than broad diversity collapse, is consistent with prior in vitro and in vivo work suggesting that antipsychotics exert a relatively selective antimicrobial pressure ^12, 37, 38^. The shared expansion of *Lactobacillus* across drug groups, alongside treatment-specific enrichment of taxa such as *Faecalibaculum, Dubosiella,* and *Akkermansia,* mirrors observations from human cohorts ^12, 14, 24^. These potential drug-specific effects on host metabolism and mucosal homeostasis warrant focus in future studies.

We found that the magnitude of drug-specific metabolomic effects exceeded that of taxonomic changes. This result suggests that metabolic remodeling may be a more sensitive readout of antipsychotic-induced microbiome disruption than compositional profiling alone. It also indicates that additional analyses focused on host metabolic effects may help resolve the contribution of host and microbiota metabolic effects of antipsychotics. Pathway enrichment analysis identified convergent disruption of lipid- and sterol-associated metabolism across all four antipsychotics, including squalene and cholesterol biosynthesis, glycerophospholipid metabolism, and cytochrome P450-associated pathways, indicating a shared class effect consistent with the known metabolic side effects of these drugs in humans ^28, 29^. Superimposed on this convergence were drug-specific signatures: haloperidol was distinguished by enrichment of sterol-to-bile acid biosynthetic pathways ^22^, olanzapine by tryptophan and amino acid catabolism, quetiapine by polyunsaturated fatty acid and porphyrin pathways, and risperidone by lipid transport and signaling.

Network-based analyses reinforced these findings at a higher organizational level. WGCNA identified coordinated metabolite modules enriched for butanoate metabolism, fatty acid activation, and redox homeostasis that were predominantly associated with vehicle-treated mice and were diminished across antipsychotic groups. The loss of these modules is notable because butyrate-associated metabolic networks are central to colonization resistance and mucosal defense ^7, 8, 33, 34^. Integration of microbiome and metabolomics data through bipartite network analysis linked specific taxa to altered metabolic states. Butyrate-associated genera such as *Butyricicoccus* and *Romboutsia* were correlated with neurometabolic and sphingolipid-related metabolites, whereas *Akkermansia* and *Staphylococcus* showed broadly inverse associations across lipid-associated pathways. Collectively, these integrated analyses suggest that antipsychotic exposure may reorganize genus-metabolite networks, with potential downstream effects on signaling environments relevant to mucosal defense and barrier integrity ^35, 36^. However, these correlations require functional validation to establish directionality and causality.

The strong positive correlation between the *Lachnospiraceae* NK4A136 group and the putatively annotated Norbadione A (*ρ* = 0.648) is notable given the established role of *Lachnospiraceae* in biotransformation of complex dietary xenobiotics ^39, 40^. This association may reflect direct degradation of diet-derived fungal or plant compounds into detectable pulvinic acid derivatives, or alternatively, given the putative nature of the annotation, the detected feature may represent a structurally analogous bacterial metabolite. The parallel positive correlations with lipid-associated metabolites such as LysoPC(16:0) and sphingosine-1-phosphate suggest that this taxon operates within a coordinated metabolic network associated with lipid homeostasis and gut barrier integrity though targeted validation is needed to confirm the identity and biological origin of this feature.

This study had notable limitations. First, all metabolite annotations from untargeted fecal metabolomic fingerprinting are putative and require confirmation through targeted MS/MS analysis with authentic standards. The biological origin of these detected features remains incompletely resolved, as fecal metabolite profiles reflect combined contributions from microbial metabolism, host metabolism, diet, xenobiotic biotransformation, and residual parent drug. Second, the integrative analyses are correlation-based and cannot establish causal relationships between specific taxa and metabolic changes. Third, this study used male mice with a single dosing regimen per drug. Sex-dependent effects and dose-response relationships remain unexplored. Finally, translation to human systems requires validation given differences in microbiome composition, diet, polypharmacy and the chronic antipsychotic exposure patterns typical of clinical use.

Together, these findings support a model in which antipsychotics disrupt gut microecology with concurrent reprogramming of microbial metabolic function rather than solely depleting microbial diversity. The convergence of these effects on pathways central to colonization resistance, including butanoate metabolism, bile acid biosynthesis, and lipid-sterol homeostasis provides a plausible mechanistic framework linking antipsychotic exposure to impaired host defense against enteric pathogens^8, 33, 41^. If validated in human cohorts, these results would position the gut microbiome as a modifiable mediator of antipsychotic-associated infection risk and suggest that microbiota-directed strategies may warrant investigation as adjunctive approaches for patients receiving antipsychotic therapy.

## Methods

### Animals

All animal procedures were approved by the Emory University Institutional Animal Care and Use Committee (IACUC; protocol number PROTO201800041) and were conducted in accordance with institutional guidelines. Conventional male specific-pathogen-free C57BL/6 mice, 6–8 weeks of age, were obtained from The Jackson Laboratory (Bar Harbor, ME) and housed under specific-pathogen-free conditions with support from the Emory Gnotobiotic Animal Core (EGAC). Mice were provided sterilized 2019 Teklad Global 19% Protein Extruded Rodent Diet (Inotiv, Indianapolis, IN) and autoclaved drinking water ad libitum. Animals were housed in hermetically sealed ISOcage P Bioexclusion units (Tecniplast, West Chester, PA) within EGAC to maintain microbiological containment. Mice were maintained under standard environmental conditions, including a 12-hour light/dark cycle, and were monitored regularly for general health, behavior, and signs of distress.

Before initiation of the experiments, microbial communities were equalized across cages through the exchange of fecal material and bedding. Each treatment group initially contained five mice, for a planned total of 25 animals. Group sizes were selected pragmatically based on the exploratory nature of the study, prior experience with the experimental models, and animal and analytical resource constraints. One mouse assigned to the quetiapine treatment group died during the bedding-exchange period before initiation of antipsychotic administration or other experimental procedures. Thus, 24 mice entered the experiment. No formal animal-level inclusion or exclusion criteria were established a priori beyond the use of apparently healthy male specific-pathogen-free mice of the specified strain and age. Assay-specific data-quality criteria were applied as described in the corresponding analytical methods.

Where indicated, antipsychotic therapeutics were administered by oral gavage. All compounds were freshly prepared in a suspension containing 0.5% (w/v) methylcellulose and 0.1% (v/v) Tween 80 in sterile water and were administered at a constant volume of 100 µL. Antipsychotics were delivered at the following doses: haloperidol, 1 mg/kg/day twice daily; olanzapine, 4 mg/kg/day twice daily; quetiapine, 100 mg/kg/day twice daily; and risperidone, 0.75 mg/kg/day twice daily, consistent with previously reported murine studies^21, 42–44^. For oral gavage, conscious mice were manually restrained using a standard scruff-restraint technique, and the dosing solution was administered using an appropriately sized gavage needle by trained personnel. Animals were not anesthetized for oral gavage because the procedure was brief and did not require surgical manipulation or other painful intervention. Avoiding anesthesia also minimized the potential for anesthetic-related effects on locomotor behavior, gastrointestinal physiology, microbial community composition, and fecal metabolomic measurements. Animals were returned to their home cages immediately after dosing and were observed for evidence of gavage-related complications or distress.

No injectable agents, including injectable anesthetics or sedatives, were administered as part of the study. Anesthesia was not used during behavioral testing, antipsychotic administration, infection monitoring, or other experimental procedures. This approach avoided the introduction of additional pharmacologic exposures that could confound behavioral, microbiological, metabolic, or infection-related outcomes.

At the designated experimental endpoint, mice were euthanized in accordance with Emory University IACUC requirements and the *AVMA Guidelines for the Euthanasia of Animals: 2020 Edition*. Euthanasia was performed by gradual-fill carbon dioxide inhalation. Carbon dioxide was introduced into the euthanasia chamber or home cage at a displacement rate of approximately 30% of the chamber or cage volume per minute to allow progressive loss of consciousness before exposure to euthanizing concentrations. Gas flow was continued until animals were unconscious and respiratory activity had ceased. Death was confirmed in accordance with the approved institutional protocol before dissection and collection of tissues, intestinal contents, and other biological samples. Carbon dioxide euthanasia was selected because it was an IACUC-approved method appropriate for mice, permitted euthanasia of animals without the administration of injectable drugs, and enabled prompt collection of biological specimens while minimizing the possibility that anesthetic or euthanasia-agent residues would alter microbiome or metabolomic measurements.

### Behavioral Testing

Open field testing was performed as previously described^45^. At the indicated time point, mice were habituated to a dedicated behavior room, in their home cages, for 1 hour. Mice were then placed in a 45cm square arena and recorded for a total of 30 minutes. Cumulative distance traveled and time spent in the center was recorded using 10 minute bins. All behavioral tracking and analysis was performed using EthoVision XT software (version 16.0.1538; Noldus Information Technology, Wageningen, the Netherlands; https://www.noldus.com/ethovision-xt) and the testing arenas were cleaned with 70% ethanol following each use.

### Citrobacter Rodentium Infection

Mice were administered antipsychotic treatments or vehicle control daily for three weeks prior to enteric challenge. *Citrobacter rodentium* (ATCC 51459) ^7, 8^ was cultured in LB medium at 37C in aerobic conditions, harvested by centrifugation at 3,000 × g for 10 min, and the bacterial pellet was resuspended in sterile Hanks’ balanced salt solution. Following the pretreatment period, mice were orally gavaged once with *C. rodentium* at a dose of 2 × 10⁹ colony-forming units (CFU). Body weight was monitored daily throughout the experiment. Fecal samples were collected at baseline and weekly following infection, homogenized, serially diluted, and plated on MacConkey agar for CFU enumeration to quantify *C. rodentium* burden.

### Computational Methods

All computational analyses were performed in RStudio using R version 4.3.3. Details of all packages, package versions, and analysis scripts are available in the publicly accessible repository hosted on Zenodo (https://doi.org/10.5281/zenodo.19337970).

### DNA Extraction Methods

Genomic DNA was extracted by SeqCenter (Pittsburgh, PA) using the ZymoBIOMICS™ DNA Miniprep Kit according to manufacturer guidelines. Briefly, samples were resuspended in lysis solution based on input type (feces), transferred to bead-beating tubes, and mechanically lysed using a FastPrep-24 system. Lysates were clarified by centrifugation, and DNA was purified using silica spin columns with sequential binding, washing, and elution steps. Eluted DNA was further cleaned using a ZymoBIOMICS™ HRC filter to remove potential inhibitors. Final DNA concentrations were quantified using a Qubit fluorometric assay.

### 16S rRNA gene amplicon sequencing and analysis

Fecal samples were analyzed by 16S rRNA gene amplicon sequencing using paired-end Illumina technology by SeqCenter (Pittsburgh, PA). Samples represented five treatment groups (Vehicle, Haloperidol, Quetiapine, Risperidone, and Olanzapine) collected longitudinally across multiple time points (T0, T2, T4, T6, and T8). Demultiplexed paired-end FASTQ files were processed using QIIME 2 (≥2022.10) 5. Primer sequences (V3/V4 Forward Trim Sequence: CCTAYGGGNBGCWGCAG; Reverse Trim Sequence: GACTACNVGGGTMTCTAATCC) were removed using the cutadapt plugin with degenerate primer matching, and sequence quality was assessed prior to denoising.

Amplicon sequence variants (ASVs) were inferred using the DADA2 plugin, which performs quality filtering, error correction, paired-end merging, and chimera removal using a consensus approach 5, 6. Taxonomic assignment was performed using a naïve Bayes classifier trained on the SILVA 138 (99% identity) reference database. Representative ASV sequences were aligned using MAFFT, hypervariable regions were masked, and a phylogenetic tree was constructed using FastTree and midpoint-rooted 5. QIIME 2 outputs were imported into R and integrated into a single phyloseq object containing the ASV abundance table, taxonomy, rooted phylogeny, and curated sample metadata 7.

Alpha diversity metrics (observed ASVs, Shannon diversity, and Simpson diversity) were calculated using phyloseq and evaluated longitudinally across time points with consistent sampling. Temporal trends were assessed using linear models with treatment-by-time interactions.

Beta diversity was assessed using Bray–Curtis dissimilarity 8. Samples were rarefied to an even sequencing depth for ordination, and principal coordinates analysis (PCoA) was used to visualize community-level differences across treatment groups and time points. Longitudinal divergence was quantified by calculating Bray–Curtis distances relative to baseline (T0).

Differential abundance testing was performed using DESeq2 with size-factor estimation by the “poscounts” method 9. Negative binomial models were used to estimate log2 fold changes, and statistical significance was assessed using Benjamini–Hochberg false discovery rate (FDR) correction. Differentially abundant taxa were visualized using volcano plots annotated at the deepest available taxonomic rank.

### High-resolution metabolomics (HRM) sample preparation

Baseline fecal samples from before onset of antipsychotic treatment and after three weeks of treatment before onset of any challenge (behavior or *C. rodentium*) was collected via clean catch and flash frozen for metabolomics analysis. Samples were prepared by the addition of 15 μl of ice-cold 33% LCMS Grade Water (Thermo Scientific 047146-K2)/66% acetonitrile-internal standard solution per milligram of sample followed by homogenization by vortexing. The homogenized samples were transferred to microcentrifuge tubes, vortexed, and placed on ice for 30 minutes before centrifugation at 14,000 × g for 10 minutes at 4 °C to precipitate the proteins. The supernatants were transferred to autosampler vials and stored at −80 °C before instrumental analysis.

### HRM Instrument Analysis

The LCMS platform consisted of a Thermo Scientific Vanquish Duo UHPLC and a Thermo Scientific Orbitrap ID-X. Two analytical platforms were used including hydrophilic interaction liquid chromatography (HILIC) coupled to positive electrospray ionization (ESI) and reversed phase C18 chromatography coupled to negative ESI. The HILIC method consisted of an Acquity BEH Amide HILIC column, 2.1 mm x 100 mm, 1.7 μm (Waters, 186004801). Buffer A was water with 1 mM ammonium acetate and 0.1% formic acid and Buffer B was 95% acetonitrile with 1 mM ammonium acetate and 0.1% formic acid. For the HILIC gradient, an initial 0.5 min hold at 90% B was followed by a linear decrease to 20% B from 0.6 to 2.55 min, a 2 min hold, and then a 5-min re-equilibration period. The C18 method consisted of a Hypersil GOLD C18 column, 2.1 x 100 mm, 1.9 μm (Thermo Scientific, 25003-102130). Buffer A was 0.1% formic acid in water and Buffer B was 99% acetonitrile with 0.1% formic acid. For the C18 gradient, an initial 0.5 min hold at 1% B was followed by a 0.75 min linear increase to 99% B, held for 3.75 min, and a 5-min re-equilibration period. Flow rates were 0.3 mL/min. The autosampler was set 4 °C and the column compartment at 45 °C. The mass spectrometer was operated at 120k resolution, and scans were collected for *m/z* 85-1,275. Tune parameters consisted of sheath gas at 50, auxiliary gas at 10, and sweep gas at 1. The spray voltage was set to 3.50 kV for ESI+ and -2.75 kV for ESI-. Ion transfer tube was set at 300 ℃ and vaporizer at 275 ℃.

### HRM data processing and statistics

Peak detection, noise removal, alignment and quantification were performed using a standardized workflow as previously described ^27^, with minor modifications as detailed: Adaptive processing for LCMS data (apLCMS) v6.3.3, with downstream quality control performed by xMSanalyzer v.2.0.8 ^19, 20^. Each metabolic feature was characterized by its *m/z* ratio, retention time, and peak intensity. For pathway enrichment analysis, differentially expressed features were normalized by log transformation, mean-centered and divided by the square root of the standard deviation of each variable and the top 10% of peaks used in *mummichog* v2.0 software ^19, 20^. Spearman correlations and heatmaps were performed and prepared using MetaboAnalyst v6.0.31 ^20^. For volcano plots, metabolomics data were imported and reshaped using the tidyverse suite in R. Intensities were normalized to z-scores within each metabolite feature. Pairwise group comparisons were performed by calculating fold changes of mean z-scores and conducting two-sided t-tests for differential abundance. Features with log2 fold change > 1 and p < 0.05 were considered significant. Volcano plots highlighting enriched and depleted metabolites were generated using ggplot2. Detected features were putatively annotated using the HMDB reference database and xMSannotator ^19, 20^.

### Weighted gene coexpression network analysis (WGCNA)

Network analyses was computed as previously described in Gacasan et al, 2025 ^27^, with minor modifications as detailed: Median-summarized, m/z-calibrated untargeted metabolomics data were processed in R (v4.3.3) using the WGCNA package (v1.72-5) ^11, 12^. Features were labeled by m/z and retention time (mz rt), log-transformed, and filtered using goodSamplesGenes to remove low-quality samples and features. Hierarchical clustering was used to assess sample relationships, and the soft-thresholding power (β = 11) was chosen using pickSoftThreshold to approximate scale-free topology. Pearson correlations were converted to a topological overlap matrix (TOM), and metabolites were clustered by average linkage on 1-TOM. Modules were detected using dynamic tree cutting (deepSplit = 2) at minClusterSize values of 100, 250, and 500 to assess granularity. A cluster size of 250 was ultimately used for downstream analysis. Module eigengenes were correlated with experimental traits (treatment groups) to identify biologically relevant modules, with significance determined by Student’s t test. Treatment groups were encoded as separate binary traits, with samples belonging to a given group coded as 1 and all remaining samples coded as 0. Thus, each module–trait correlation represents the association between a module eigengene and membership in that treatment group relative to all other groups combined; no single baseline or reference group was used. Module assignments, eigengenes, correlations, and annotated feature lists were exported for downstream interpretation.

#### Untargeted metabolomics preprocessing

Metabolomics feature tables were imported using readr and processed with dplyr/tidyr. Samples were intersected with the 16S dataset, and all matrices were reordered to identical SampleID order (verified by strict checks). Metabolites with >20% missing values were removed; remaining missing values were imputed using a half-minimum approach. Feature intensities were log-transformed (log1p) and autoscaled (z-score per feature).

### Microbe–Metabolite Integration Overview

Integration was performed at a single pre-challenge time point (Week 3) to ensure matched samples across assays and to avoid longitudinal non-independence. The 16S rRNA gene sequencing data were managed in phyloseq and subset to Week 3, with zero-abundance taxa removed. Sample metadata were extracted with SampleID retained explicitly to enforce alignment across data modalities. Microbiome data were transformed using centered log-ratio (CLR) transformation after appropriate filtering, and metabolite intensities were log-transformed and filtered based on variance. Pairwise associations between microbial taxa and metabolites were assessed using Spearman rank correlation.

### Spearman Correlation Heatmap

Spearman correlation coefficients (ρ) were calculated between selected microbial taxa (e.g., genus level) and metabolite features across matched samples. Features were filtered based on prevalence and variance prior to analysis. Correlation matrices were visualized as clustered heatmaps using hierarchical clustering (Euclidean distance, Ward’s method), with color indicating correlation strength and direction.

### Bipartite Network Analysis

Significant microbe–metabolite associations were used to construct a bipartite network. Nodes represented microbial taxa and metabolites, and edges represented significant correlations exceeding a predefined threshold (e.g., |ρ| ≥ 0.4). Edge thickness was scaled to correlation magnitude, and sign was encoded by color. Network visualization and layout were performed using **igraph** or **mixOmics**-based functions.

### Circos Plot Visualization

To visualize multi-omics associations, circos plots were generated using the **mixOmics** DIABLO framework. Features selected by the model were included based on their contribution to component separation and cross-block correlation. Associations exceeding a specified correlation threshold were displayed as links between microbial and metabolite features. Outer tracks represent feature loadings or group-level expression patterns, while inner connections represent pairwise correlations.

#### Visualization and network analysis

Integrated associations were visualized using heatmaps generated with ComplexHeatmap and circlize, and as bipartite correlation networks constructed with igraph and rendered using ggraph/tidygraph. Repelled node labels were added with ggrepel, and plots were styled using ggplot2. Targeted subnetworks highlighted genera of interest (e.g., *Akkermansia*, Lachnospiraceae-associated taxa, *Staphylococcus*) and their directly connected metabolite neighborhoods.

All analyses were performed in R using reproducible, script-based workflows, with intermediate tables and figures exported for downstream analysis and figure assembly.

### Statistics

Statistical analysis was performed using Prism 10 (GraphPad, San Diego, CA). Significance is defined as *P ≤ .05, **P ≤ .01, ***P ≤ .001, ****P ≤ .0001

## Data Availability Statement

Raw metabolomics data generated in this study are available through Metabolomics Workbench (study DOIs pending). Untargeted metabolomics feature tables and additional datasets are available via the Emory Dataverse.

Sequencing data have been deposited in the NCBI Sequence Read Archive under BioProject accession **PRJNA1445182** at https://www.ncbi.nlm.nih.gov/bioproject/1445182.

## Code Availability

All computational code used for microbiome profiling, metabolomics analysis, and integrative multiomic workflows is publicly available via Zenodo at https://doi.org/10.5281/zenodo.19337970. This repository includes scripts for 16S rRNA analysis, statistical modeling, data visualization, and multiomic integration, including correlation and network based approaches.

All other relevant data supporting the findings of this study are available from the corresponding author upon reasonable request.

## Author Contributions

CAG, RMJ, TRS, and MW conceived and designed the experiments. LDL and TRS conducted murine open field testing. CAG conducted all other mouse experiments. CAG conducted sample preparation for LC-HRMS. Mass spectrometry data acquisition was performed by JW and DPJ. All downstream HRMS analysis was performed by CAG. All other analyses and experiments were conducted and analyzed by CAG and GW. CAG, RMJ, and MW wrote the manuscript.

## Acknowledgments

The Emory Integrated Cellular Imaging Core (RRID:SCR_023534) and the Emory Cancer Tissue and Pathology Shared Resource are subsidized by the Emory University School of Medicine. The Emory Gnotobiotic Animal Core especially Caroline Addis and Amanda Metzger for their assistance with animal handling as well as Sean D. Kelly from the Sampson Lab for dissection assistance.

This study was funded by a NRSA F30 training grant from the National Institute of Diabetes and Digestive and Kidney Diseases [*F30 DK139762*], and the Emory Prevention Epicenter Program (PEACH) through the Centers for Disease Control and Prevention (CDC) [*U54CK000601*] as well as the National Institute for Allergy and Infectious Diseases [*K23AI144036*] and in part by an Imagine, Innovate and Impact (I3) from the Emory School of Medicine and through the Georgia CTSA NIH award (*UL1-TR002378*). The funder played no role in study design, data collection, analysis and interpretation of data, or the writing of this manuscript. This content is solely the responsibility of the authors and does not necessarily represent the official view of the National Institutes of Health or Centers for Disease Control and Prevention.

