## Supplementary for "Anti-psychotic therapeutics modify gut microbial metabolism and modulate susceptibility to gastrointestinal infection in mice"

**SUPPLEMENTARY DATA**

C. Anthony Gacasan[1], Jaclyn Weinberg[2], Lyndsey D. Lipson[3], Gabrielle Webster[1], Dean P. Jones[2], Timothy Sampson[3], Rheinallt Jones[1], Michael H. Woodworth[4*]

Affiliations:

[1]Division of Gastroenterology, Hepatology and Nutrition Department of Pediatrics, Emory University School of Medicine, Atlanta, GA, USA.

[2]Division of Pulmonary, Allergy, Critical Care and Sleep Medicine, Department of Medicine, Emory University School of Medicine, Atlanta, GA

[3]Department of Cell Biology, Emory University School of Medicine, Atlanta, GA 30322, United States

[4]Division of Infectious Disease, Department of Medicine, Emory University School of Medicine, Atlanta, GA, USA

Key words: Anti-Psychotics, Microbiome

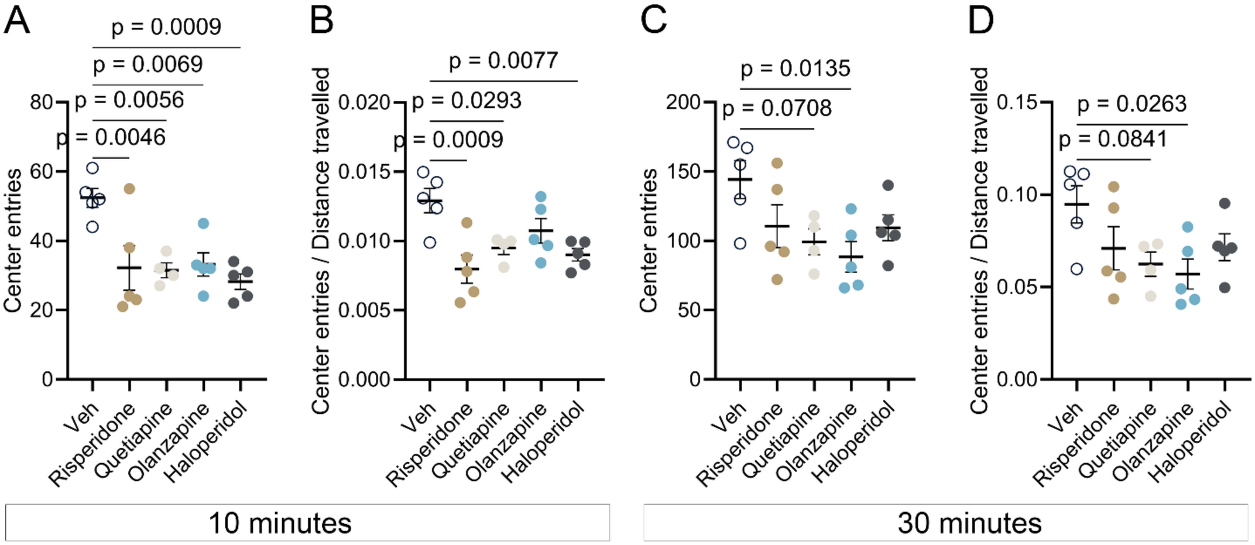


**Supplementary Figure 1.** Center-entry behavior during the open-field test. (A) Number of center entries during the first 10 minutes of the open-field test. (B) Center entries during the first 10 minutes normalized to distance traveled. (C) Number of center entries during the full 30-minute observation period. (D) Center entries during the 30-minute observation period normalized to distance traveled. Data are presented as mean ± SEM. P values were calculated using one-way ANOVA.

| contrast | estimate | Standard Error | Degrees of Freedom | t-Statistic | p-value | Adjusted p-value (FDR) |
|---|---|---|---|---|---|---|
| *Risperidone - Vehicle* | 0.26297 | 0.06952 | 62 | 3.782649 | 0.000352 | 0.001407 |
| *Olanzapine - Vehicle* | 0.142311 | 0.071972 | 62 | 1.977294 | 0.052459 | 0.157377 |
| *Haloperidol - Vehicle* | 0.101943 | 0.06952 | 62 | 1.466388 | 0.147597 | 0.295193 |
| *Quetiapine - Vehicle* | 0.023315 | 0.071145 | 62 | 0.327706 | 0.744238 | 0.744238 |

**Supplementary Table 1.** ANCOVA analysis of *C. rodentium* colonization dynamics over time. $Log_{10}$-transformed stool bacterial burden (CFU per gram of stool) was modeled as a function of time and treatment group using analysis of covariance (ANCOVA) to assess differences in temporal slopes between antipsychotic treatment groups and vehicle control. Reported statistics summarize treatment-specific slope estimates and associated significance testing, with multiple-comparison adjustment applied where indicated.

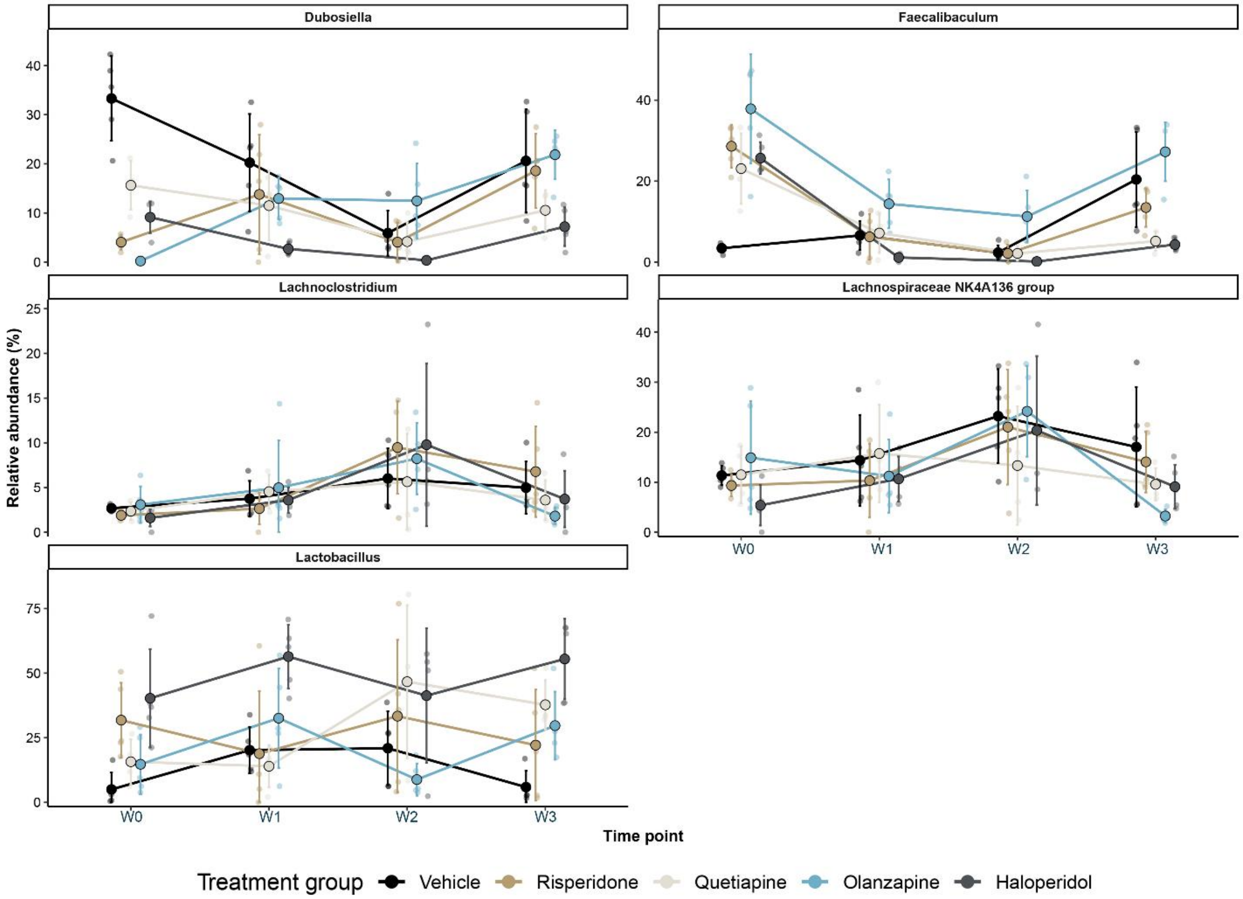


**Supplementary Figure 2. Longitudinal changes in the relative abundance of dominant bacterial genera following antipsychotic treatment.** Relative abundances of Faecalibaculum, Dubosiella, Lachnospiraceae NK4A136 group, Lachnoclostridium, and Lactobacillus are shown across W0, W1, W2 and W3 for vehicle-, risperidone-, quetiapine-, olanzapine-, and haloperidol-treated mice. Small points represent individual mice, large points represent group means, and error bars indicate mean ± standard deviation. Lines connect group means across time and are provided to visualize treatment-specific temporal patterns. Temporal trajectories were compared using generalized estimating equations with categorical time, treatment group, and their interaction, with mouse specified as the repeated-measures cluster and an AR(1) working correlation structure. Relative abundances were arcsine-square-root transformed before statistical analysis. Pairwise whole-trajectory Wald tests jointly compared between-group changes from baseline at (W0) at W1,

W2 and W3, with P values corrected within each taxon using the Benjamini–Hochberg false discovery rate procedure. Complete pairwise statistical comparisons are provided in Supplementary Table X.

| Taxon | Group_comparison | df1 | df2 | F.ratio | p.value | Wald_chisq | Wald_df | p_FDR | Significance |
|---|---|---|---|---|---|---|---|---|---|
| Faecalibaculum | *Vehicle - Risperidone* | *3* | *75* | *44.073* | *1.58E-16* | *132.219* | *3* | *7.89E-16* | **** |
| Faecalibaculum | *Vehicle - Quetiapine* | *3* | *75* | *33.989* | *5.58E-14* | *101.967* | *3* | *1.86E-13* | **** |
| Faecalibaculum | *Vehicle - Olanzapine* | *3* | *75* | *17.905* | *7.33E-09* | *53.715* | *3* | *1.83E-08* | **** |
| Faecalibaculum | *Vehicle - Haloperidol* | *3* | *75* | *69.542* | *1.31E-21* | *208.626* | *3* | *1.31E-20* | **** |
| Faecalibaculum | *Risperidone - Quetiapine* | *3* | *75* | *2.689* | *0.052333* | *8.067* | *3* | *0.073574* | *ns* |
| Faecalibaculum | *Risperidone - Olanzapine* | *3* | *75* | *0.806* | *0.494417* | *2.418* | *3* | *0.494417* | *ns* |
| Faecalibaculum | *Risperidone - Haloperidol* | *3* | *75* | *8.498* | *6.26E-05* | *25.494* | *3* | *1.25E-04* | *** |
| Faecalibaculum | *Quetiapine - Olanzapine* | *3* | *75* | *1.925* | *0.132834* | *5.775* | *3* | *0.147593* | *ns* |
| Faecalibaculum | *Quetiapine - Haloperidol* | *3* | *75* | *3.367* | *0.022935* | *10.101* | *3* | *0.038226* | * |
| Faecalibaculum | *Olanzapine - Haloperidol* | *3* | *75* | *2.593* | *0.058859* | *7.779* | *3* | *0.073574* | *ns* |
| Dubosiella | *Vehicle - Risperidone* | *3* | *75* | *14.98* | *9.85E-08* | *44.94* | *3* | *1.64E-07* | **** |
| Dubosiella | *Vehicle - Quetiapine* | *3* | *75* | *3.45* | *0.020718* | *10.35* | *3* | *0.02302* | * |
| Dubosiella | *Vehicle - Olanzapine* | *3* | *75* | *35.16* | *2.69E-14* | *105.48* | *3* | *6.73E-14* | **** |
| Dubosiella | *Vehicle - Haloperidol* | *3* | *75* | *4.878* | *0.003748* | *14.634* | *3* | *0.004685* | ** |
| Dubosiella | *Risperidone - Quetiapine* | *3* | *75* | *35.434* | *2.27E-14* | *106.302* | *3* | *6.73E-14* | **** |
| Dubosiella | *Risperidone - Olanzapine* | *3* | *75* | *18.607* | *4.03E-09* | *55.821* | *3* | *8.07E-09* | **** |
| Dubosiella | *Risperidone - Haloperidol* | *3* | *75* | *13.825* | *2.89E-07* | *41.475* | *3* | *4.12E-07* | **** |
| Dubosiella | *Quetiapine - Olanzapine* | *3* | *75* | *67.093* | *3.48E-21* | *201.279* | *3* | *3.48E-20* | **** |
| Dubosiella | *Quetiapine - Haloperidol* | *3* | *75* | *0.447* | *0.720114* | *1.341* | *3* | *0.720114* | *ns* |
| Dubosiella | *Olanzapine - Haloperidol* | *3* | *75* | *50.373* | *6.11E-18* | *151.119* | *3* | *3.06E-17* | **** |
| Lachnospiraceae NK4A136 group | *Vehicle - Risperidone* | *3* | *75* | *0.316* | *0.813969* | *0.948* | *3* | *0.813969* | *ns* |
| Lachnospiraceae NK4A136 group | *Vehicle - Quetiapine* | *3* | *75* | *2.506* | *0.065415* | *7.518* | *3* | *0.108321* | *ns* |
| Lachnospiraceae NK4A136 group | *Vehicle - Olanzapine* | *3* | *75* | *3.709* | *0.015147* | *11.127* | *3* | *0.050491* | *ns* |
| Lachnospiraceae NK4A136 group | *Vehicle - Haloperidol* | *3* | *75* | *0.882* | *0.454399* | *2.646* | *3* | *0.504887* | *ns* |
| Lachnospiraceae NK4A136 group | *Risperidone - Quetiapine* | *3* | *75* | *3.208* | *0.027827* | *9.624* | *3* | *0.069567* | *ns* |
| Lachnospiraceae NK4A136 group | *Risperidone - Olanzapine* | *3* | *75* | *14.924* | *1.04E-07* | *44.772* | *3* | *1.04E-06* | **** |
| Lachnospiraceae NK4A136 group | *Risperidone - Haloperidol* | *3* | *75* | *2.385* | *0.075825* | *7.155* | *3* | *0.108321* | *ns* |
| Lachnospiraceae NK4A136 group | *Quetiapine - Olanzapine* | *3* | *75* | *2.938* | *0.038642* | *8.814* | *3* | *0.077285* | *ns* |
| Lachnospiraceae NK4A136 group | *Quetiapine - Haloperidol* | *3* | *75* | *2.073* | *0.110874* | *6.219* | *3* | *0.138593* | *ns* |
| Lachnospiraceae NK4A136 group | *Olanzapine - Haloperidol* | *3* | *75* | *6.199* | *8.07E-04* | *18.597* | *3* | *0.004033* | ** |
| Lachnoclostridium | *Vehicle - Risperidone* | *3* | *75* | *2.277* | *0.086498* | *6.831* | *3* | *0.172996* | *ns* |
| Lachnoclostridium | *Vehicle - Quetiapine* | *3* | *75* | *1.053* | *0.374408* | *3.159* | *3* | *0.451601* | *ns* |
| Lachnoclostridium | *Vehicle - Olanzapine* | *3* | *75* | *4.404* | *0.006582* | *13.212* | *3* | *0.032909* | * |
| Lachnoclostridium | *Vehicle - Haloperidol* | *3* | *75* | *1.659* | *0.18303* | *4.977* | *3* | *0.305051* | *ns* |
| Lachnoclostridium | *Risperidone - Quetiapine* | *3* | *75* | *3.078* | *0.032567* | *9.234* | *3* | *0.108556* | *ns* |
| Lachnoclostridium | *Risperidone - Olanzapine* | *3* | *75* | *7.083* | *2.96E-04* | *21.249* | *3* | *0.002964* | ** |

| | | | | | | | | | |
|---|---|---|---|---|---|---|---|---|---|
| Lachnoclostridium | *Risperidone - Haloperidol* | *3* | *75* | *2.658* | *0.054356* | *7.974* | *3* | *0.13589* | *ns* |
| Lachnoclostridium | *Quetiapine - Olanzapine* | *3* | *75* | *0.888* | *0.451601* | *2.664* | *3* | *0.451601* | *ns* |
| Lachnoclostridium | *Quetiapine - Haloperidol* | *3* | *75* | *0.955* | *0.418469* | *2.865* | *3* | *0.451601* | *ns* |
| Lachnoclostridium | *Olanzapine - Haloperidol* | *3* | *75* | *0.914* | *0.438609* | *2.742* | *3* | *0.451601* | *ns* |
| Lactobacillus | *Vehicle - Risperidone* | *3* | *75* | *4.387* | *0.006716* | *13.161* | *3* | *0.009594* | ** |
| Lactobacillus | *Vehicle - Quetiapine* | *3* | *75* | *38.991* | *2.71E-15* | *116.973* | *3* | *2.71E-14* | **** |
| Lactobacillus | *Vehicle - Olanzapine* | *3* | *75* | *8.82* | *4.42E-05* | *26.46* | *3* | *1.47E-04* | *** |
| Lactobacillus | *Vehicle - Haloperidol* | *3* | *75* | *5.2* | *0.002566* | *15.6* | *3* | *0.004277* | ** |
| Lactobacillus | *Risperidone - Quetiapine* | *3* | *75* | *10.648* | *6.54E-06* | *31.944* | *3* | *3.27E-05* | **** |
| Lactobacillus | *Risperidone - Olanzapine* | *3* | *75* | *6.698* | *4.57E-04* | *20.094* | *3* | *0.001143* | ** |
| Lactobacillus | *Risperidone - Haloperidol* | *3* | *75* | *2.648* | *0.055036* | *7.944* | *3* | *0.061151* | *ns* |
| Lactobacillus | *Quetiapine - Olanzapine* | *3* | *75* | *6.093* | *9.10E-04* | *18.279* | *3* | *0.001821* | ** |
| Lactobacillus | *Quetiapine - Haloperidol* | *3* | *75* | *3.128* | *0.030657* | *9.384* | *3* | *0.038321* | * |
| Lactobacillus | *Olanzapine - Haloperidol* | *3* | *75* | *0.108* | *0.955236* | *0.324* | *3* | *0.955236* | *ns* |

**Supplementary Table 2. Pairwise comparisons of longitudinal relative-abundance trajectories among treatment groups for selected bacterial genera.** Temporal trajectories of *Faecalibaculum*, *Dubosiella*, Lachnospiraceae NK4A136 group, *Lachnoclostridium*, and *Lactobacillus* were compared using generalized estimating equations with treatment group, categorical time, and their interaction as fixed effects and mouse as the repeated-measures cluster. Relative abundances were arcsine-square-root transformed before analysis. Wald tests jointly evaluated between-group differences in changes from baseline (W0) at W1, W2, and W3. The Benjamini–Hochberg procedure was applied to the 10 pairwise comparisons within each genus. Wald $\chi^2$ tests had 3 degrees of freedom. Statistical significance was defined as BH-FDR $q < 0.05$. *$q < 0.05$; **$q < 0.01$; ***$q < 0.001$; ****$q < 0.0001$; ns, not significant.

| | F.Model | R2 | p-value | Adjusted p-value (FDR) |
|---|---|---|---|---|
| *H_Post vs O_Post* | 7.1388 | 0.50491 | 0.028 | 0.035 |
| *H_Post vs Q_Post* | 16.182 | 0.69804 | 0.007 | 0.026 |
| *H_Post vs R_Post* | 1.6586 | 0.17172 | 0.229 | 0.229 |
| *H_Post vs V_Post* | 17.376 | 0.68475 | 0.007 | 0.026 |
| *O_Post vs Q_Post* | 19.089 | 0.76085 | 0.025 | 0.035 |
| *O_Post vs R_Post* | 7.5195 | 0.51789 | 0.02 | 0.033333 |
| *O_Post vs V_Post* | 3.8043 | 0.35211 | 0.032 | 0.035556 |
| *Q_Post vs R_Post* | 8.0274 | 0.53418 | 0.012 | 0.026 |
| *Q_Post vs V_Post* | 34.829 | 0.83265 | 0.008 | 0.026 |
| *R_Post vs V_Post* | 14.923 | 0.651 | 0.013 | 0.026 |

**Supplementary Table 3.** Pairwise permutational multivariate analysis of variance (PERMANOVA) results for untargeted metabolomics profiles across post-treatment groups. PERMANOVA was performed using all measured metabolomic features to assess differences in overall metabolic composition between each pair of treatment groups at the post-treatment time point. Reported statistics include the pseudo-F statistic (F.Model), proportion of variance explained ($R^2$), nominal p-values (pval), and multiple-testing–adjusted p-values (p.adj). Adjusted p-values were calculated to account for multiple pairwise comparisons.

| Name | mz_rt | Haloperidol Average Intensity | Olanzapine Average Intensity | Quetiapine Average Intensity | Risperidone Average Intensity | Vehicle Average Intensity | p_adj_fdr | HMDB Class | HMDB Sub Class |
|---|---|---|---|---|---|---|---|---|---|
| *Nornicotine* | 149.107318_104.3185079 | 26104574.8 | 39864430 | 11310177 | 24966371 | 62027410.4 | 1.98E-07 | Pyridines and derivatives | |
| *Diethylpropion* | 206.1538863_98.83902698 | 934757 | 1815774 | 537459.3 | 890646.2 | 2903758.2 | 2.93E-07 | Organo ocygen compounds | |
| *N-Feruloylglycyl-L-phenylalanine* | 399.155949_79.73455007 | 1203151 | 1658022 | 30950782 | 1043441.6 | 1256608.4 | 2.93E-07 | Carboxylic acids and derivatives | Amino acids, peptides, and analogues |
| *Castamollissin* | 469.0994193_150.3065675 | 0 | 0 | 223207.5 | 0 | 0 | 2.93E-07 | Organooxygen compounds | Carbohydrates and carbohydrate conjugates |
| *Phenelzine* | 137.1072823_97.18838279 | 8731314.4 | 15285975 | 4160314 | 8300971 | 18648463.4 | 3.21E-07 | Benzene and substituted derivatives | Not Available |
| *Myricetin 33-digalactoside* | 643.1503175_45.22017116 | 7515757 | 19275838 | 5585051 | 7312142 | 25523894.8 | 5.09E-07 | Flavonoids | Flavonoid glycosides |
| *Agrocybenine* | 207.1492354_100.4989209 | 16713295.6 | 34227483 | 8898023 | 15640620 | 43263666.8 | 6.18E-07 | Pyrrolopyridines | Not Available |
| *Arecoline* | 156.101941_288.0266019 | 455927.6 | 1444199 | 787487.8 | 816838.4 | 1681520.8 | 8.47E-07 | Alkaloids and derivatives (SuperClass) | Not Available |
| *Crotamiton* | 204.1383012_94.52925532 | 572902.6 | 1071530 | 248828 | 532272 | 2061861 | 1.20E-06 | Benzene and substituted derivatives | Anilides |
| *Lidocaine* | 235.1803785_93.62902346 | 17994288.6 | 37363840 | 10450742 | 17988323 | 52902968.4 | 1.20E-06 | Benzene and substituted derivatives | Xylenes |
| *19-Hydroxy-PGE2* | 351.2163006_56.82984427 | 4654404 | 6358501 | 2930249 | 3785197.2 | 7731277.2 | 1.64E-06 | Fatty Acyls | Eicosanoids |
| *Petunidin 3-galactoside* | 480.1247261_74.51755109 | 0 | 0 | 337577.5 | 0 | 0 | 1.64E-06 | Flavonoids | Flavonoid glycosides |
| *Gomphrenin II* | 697.1855184_45.46218913 | 616455.2 | 1254885 | 403038.8 | 549719.2 | 1917658.8 | 1.64E-06 | Organooxygen compounds | Carbohydrates and carbohydrate conjugates |
| *(-)-3-(2-methyl-3-furyl)thio-2-butanone* | 185.0632845_116.5893885 | 1762651 | 2094326 | 748843.5 | 1411016.8 | 2588423.8 | 1.84E-06 | Thioethers | Aryl thioethers |
| *(EE)-Piperlonguminine* | 274.1436673_94.12987461 | 1193384.8 | 2528360 | 423960.5 | 976559.8 | 3728121 | 1.84E-06 | Benzodioxoles | Not Available |
| *(Cyclohexylmethyl)pyrazine* | 177.1385509_89.34431704 | 21041414 | 50973701 | 13362560 | 24156460 | 80332087.6 | 2.29E-06 | Diazines | Pyrazines |
| *L(-)-Nicotine pestanal* | 163.1229478_97.20241098 | 11617332.6 | 20794116 | 5673837 | 12049022 | 29247137.2 | 2.73E-06 | Pyridines and derivatives | Pyrrolidinylpyridines |
| *2-Methoxy-3-methyl-9H-carbazole* | 212.1070117_92.6543102 | 505652.2 | 887951.8 | 333583.5 | 525482 | 1287828.6 | 2.81E-06 | Indoles and derivatives | Carbazoles |
| *L-Phenylalanine* | 166.0861563_120.5481476 | 10288863.2 | 11997062 | 5547659 | 9265538 | 15445977 | 3.22E-06 | Carboxylic acids and derivatives | Amino acids, peptides, and analogues |
| *DG(18:3(6Z9Z12Z)22:6(4Z7Z10Z13Z16Z19Z)0:0)* | 663.4964569_45.74883329 | 464743 | 1192361 | 328251 | 478695.2 | 1580755.4 | 3.22E-06 | Glycerolipids | Diradylglycerols |
| *2367-Tetrahydro-7-methylcyclopentbazepin-8(1H)-one* | 164.106993_104.1648628 | 19434048.6 | 27230408 | 7568724 | 17033627 | 37974176.6 | 4.03E-06 | Not Available | Not Available |
| *Mexiletine* | 180.1382658_111.982739 | 944286.2 | 1629348 | 551408.5 | 819103.8 | 1815107 | 4.10E-06 | Phenol ethers | Not Available |
| *Gadoversetamide* | 663.1595836_45.42759137 | 533219.2 | 1296175 | 427680.5 | 558787.4 | 1693204.6 | 4.30E-06 | Not Available | Not Available |
| *5678-Tetrahydro-24-dimethylquinoline* | 162.1276513_87.40353705 | 1066207.4 | 1957429 | 615456.3 | 1228821.2 | 3044595 | 4.56E-06 | Quinolines and derivatives | Hydroquinolines |
| *PG(18:018:0)* | 779.5786924_45.04181284 | 641086.6 | 1616777 | 574249.5 | 699685.2 | 2416890.6 | 4.60E-06 | Glycerophospholipids | Glycerophosphoglycerols |

| | | | | | | | | | |
|---|---|---|---|---|---|---|---|---|---|
| *NO-Didesmethyltramadol* | 236.1643497_101.6198847 | 1532236.8 | 2272254 | 807111.5 | 1549279.2 | 3402785.6 | 5.10E-06 | Benzene and substituted derivatives | Cyclohexylphenols |
| *4alpha-Formyl-4beta-methyl-5alpha-cholesta-824-dien-3beta-ol* | 409.3458408_47.75421323 | 512499.4 | 64465 | 653962.3 | 674579.8 | 0 | 5.92E-06 | Prenol lipids | Triterpenoids |
| *Tocainide* | 193.1335221_111.8710709 | 177427471.6 | 3.03E+08 | 86962336 | 151571742 | 379300067.2 | 5.92E-06 | Benzene and substituted derivatives | Xylenes |
| *PC(20:5(5Z8Z11Z14Z17Z)24:1(15Z))* | 890.6642249_45.78636381 | 351826.6 | 657073.8 | 222051.8 | 311565.8 | 896298.8 | 6.66E-06 | Glycerophospholipids | Glycerophosphocholines |
| *4-Amino-2-methyl-1-naphthol* | 174.091306_99.655045 | 1161259.2 | 2234088 | 784642.8 | 1258786 | 2806615.2 | 6.68E-06 | | |
| *Kynuramine* | 165.1022653_127.5526863 | 31566857.6 | 45237190 | 14751656 | 33624749 | 62979039.8 | 7.53E-06 | Organooxygen compounds | Carbonyl compounds |
| *Cholesteryl acetate* | 429.3726654_47.83884678 | 32296665.2 | 749692.3 | 39444760 | 36352145 | 42914.8 | 8.24E-06 | Steroids and steroid derivatives | Steroid esters |
| *Tetrahydroharmol* | 203.1179045_116.1418549 | 5521338 | 8325396 | 2499932 | 5119269.2 | 11084577.6 | 8.24E-06 | Harmala alkaloids | Not Available |
| *N-Cyclopropyl-trans-2-cis-6-nonadienamide* | 176.1432981_80.76500173 | 1090134.2 | 1921665 | 523498.8 | 1561413.6 | 3849442.6 | 1.16E-05 | Fatty Acyls | Fatty amides |
| *1-Cyano-2-hydroxy-3-butene* | 98.06005797_36.87901628 | 173814.4 | 282871.3 | 224931.8 | 187072.8 | 282022.8 | 1.38E-05 | Organooxygen compounds | Alcohols and polyols |
| *lysoPC(26:1(5Z))* | 634.4781275_46.36550682 | 258989.6 | 562329.8 | 147100.5 | 238720.6 | 738681.8 | 1.72E-05 | Glycerophospholipids | Glycerophosphocholines |
| *Dictyoquinazol A* | 313.1192955_104.7398639 | 885889.6 | 804533.3 | 4859186 | 713113 | 875143 | 1.72E-05 | Diazanaphthalenes | Benzodiazines |
| *2-hydroxymexiletine* | 196.1332089_105.4920339 | 13250047.2 | 20776773 | 6640363 | 11378178 | 23769388.6 | 1.72E-05 | Phenols | 4-alkoxyphenols |
| *Metanephrine* | 198.1124197_143.5468325 | 6471391 | 8093136 | 2970160 | 5442955.8 | 8546176.4 | 1.74E-05 | Phenols | Methoxyphenols |
| *91013-Trihydroxystearic acid* | 333.2635902_57.75068644 | 12926192.4 | 15357533 | 4692815 | 7843860.6 | 19592390.2 | 1.77E-05 | Fatty Acyls | Fatty acids and conjugates |
| *Hexylcaine* | 262.1799698_90.46435654 | 1214866 | 1308012 | 467853 | 1313789.4 | 2776608.4 | 1.81E-05 | Benzene and substituted derivatives | Benzoic acids and derivatives |
| *Cyclopentolate* | 292.1906435_91.73460507 | 1123363.2 | 1426687 | 466662 | 1167837 | 2215912.2 | 2.10E-05 | Benzene and substituted derivatives | Not Available |
| *Spinosin C* | 755.2180662_45.96330447 | 354906.4 | 781138.3 | 315200.8 | 346776.6 | 1166125 | 2.18E-05 | Flavonoids | Flavonoid glycosides |
| *1-(23-Dihydro-1H-pyrrolizin-5-yl)-14-pentanedione* | 206.1175322_102.5465388 | 12080714 | 16500091 | 5942034 | 13453923 | 26867961.2 | 2.22E-05 | Carboxylic acids and derivatives | Amino acids, peptides, and analogues |
| *Nizatidine* | 332.1225878_3.746346135 | 0 | 0 | 122641.9 | 0 | 0 | 2.26E-05 | Azoles | Thiazoles |
| *Mepivacaine* | 247.180474_90.77703602 | 2590125.8 | 5316368 | 1684918 | 2966797 | 7463562.8 | 2.26E-05 | Piperidines | Piperidinecarboxylic acids and derivatives |
| *S-Ethyl thioacetate* | 105.0368462_174.2860294 | 404522.8 | 808212.3 | 198007.5 | 522394.2 | 1260368 | 2.30E-05 | Thiocarboxylic acids and derivatives | Thioesters |
| *5678-Tetrahydroquinoxaline* | 135.0916016_117.651223 | 34163326.2 | 45819091 | 15141099 | 30633181 | 58161786.8 | 2.30E-05 | Diazines | Pyrazines |
| *5-Methoxydimethyltryptamine* | 219.1491763_96.41654362 | 7790326.6 | 13380669 | 4401435 | 8241204.6 | 19180730 | 3.05E-05 | Indoles and derivatives | Tryptamines and derivatives |
| *Prostaglandin E2* | 353.2319464_57.20320237 | 5194545 | 6757572 | 2787330 | 3880331.4 | 8136562.2 | 3.69E-05 | Fatty Acyls | Eicosanoids |

**Supplementary Table 4.** Top 50 putatively annotated metabolites identified from untargeted metabolomics analysis using xMSannotator. Metabolites are ranked by false discovery rate–adjusted p-values (p_adj_fdr) derived from differential abundance testing across antipsychotic treatment groups and vehicle. For each feature, the table reports the putative compound name, m/z–retention time identifier (mz_rt), average feature intensity within each treatment group, FDR-adjusted p-value, and Human Metabolome Database (HMDB) chemical class and subclass where available. Annotations are putative and based on accurate mass, retention time, and database-supported chemical inference.

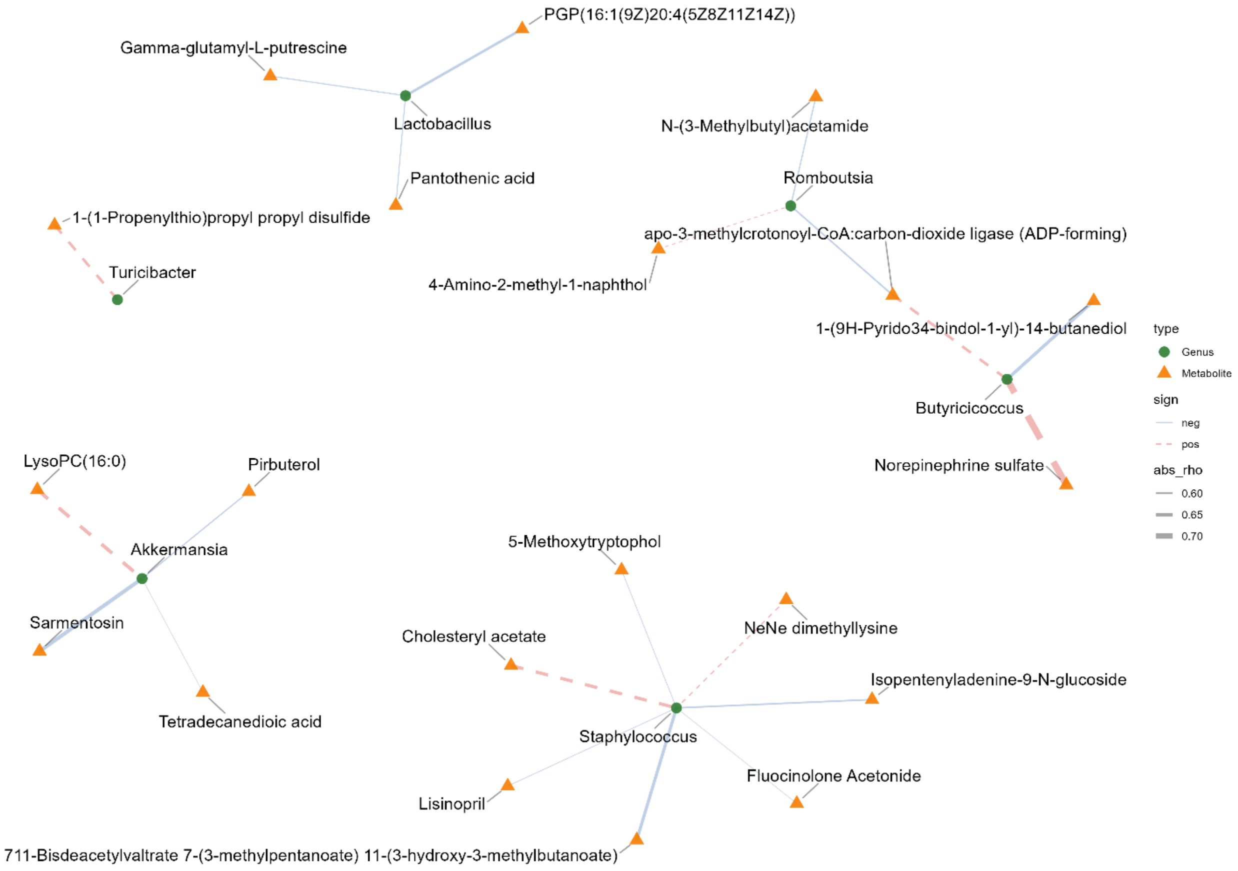


**Supplementary Figure 3.** Selected hub genera from the bipartite genus–metabolite association network constructed using the strongest correlations for visualization. Nodes represent genera (green circles) and metabolites (orange triangles). Edges are weighted by the absolute value of the correlation coefficient (|ρ|; thicker edges indicate stronger associations) and are colored and styled by directionality (positive = red dashed; negative = blue solid). Highlighted genera include *Akkermansia*, *Lactobacillus*, *Turicibacter*, *Butyricicoccus*, *Staphylococcus*, and *Romboutsia*, along with their first-degree (1-hop) neighboring metabolites.

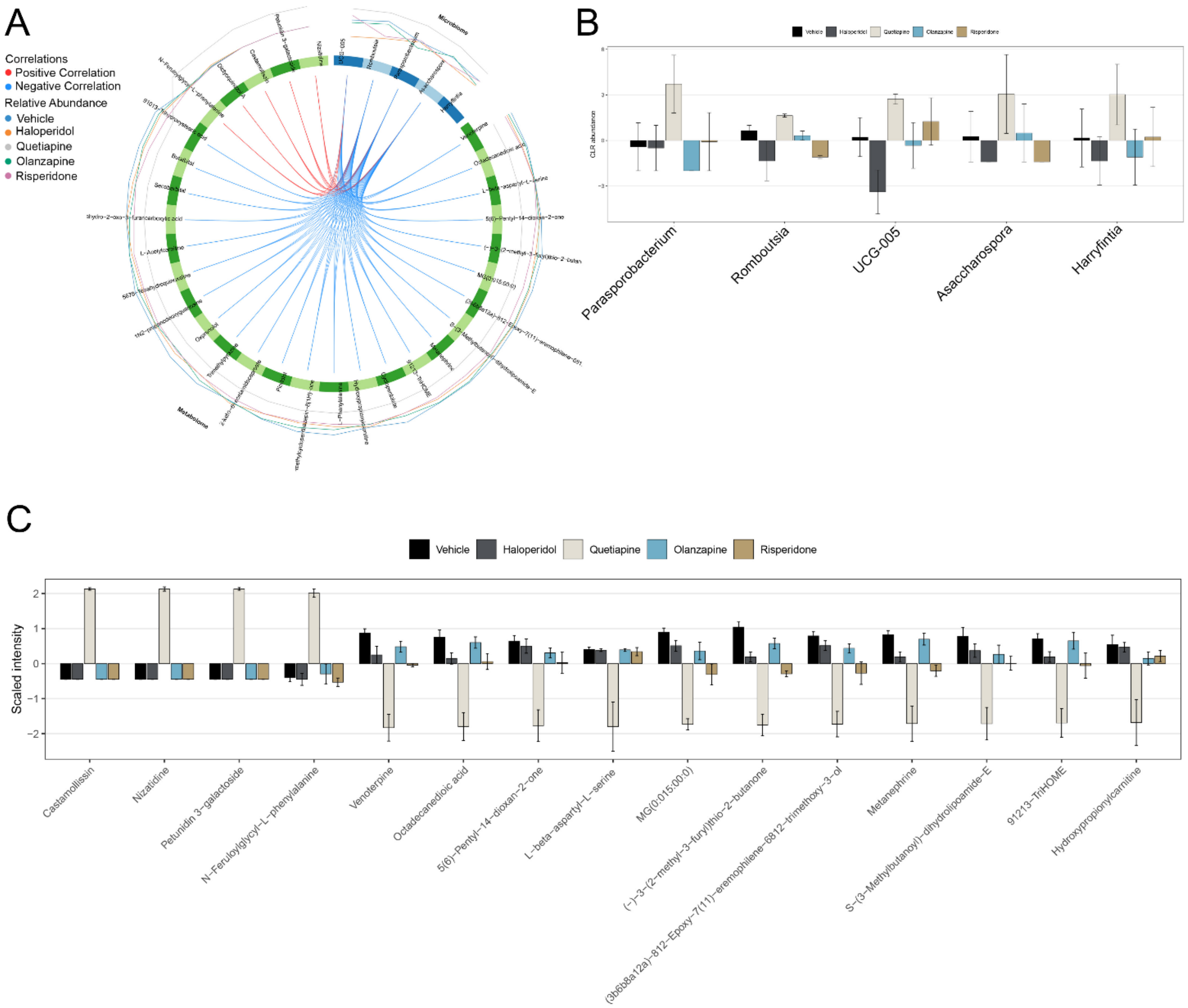


**Supplementary Figure 4. DIABLO integration of microbiome and metabolome features at week 3. (A)** Circos plot showing genus-level microbial features and putatively annotated metabolites selected by the DIABLO block.sPLS-DA model at T6. Outer tracks represent the relative contribution of each selected feature to treatment-group discrimination, and connecting lines indicate cross-omics correlations on component 1 with $|r| \geq 0.4$. Red lines indicate positive correlations and blue lines indicate negative correlations. **(B)** CLR-transformed abundances of the selected bacterial genera across treatment groups. **(C)** Scaled intensities of the selected putatively annotated metabolite features across treatment groups. Bars represent group means with error bars indicating SEM. Microbiome data were CLR-transformed following prevalence and variance filtering. Metabolomics data were log2-transformed, imputed using half of the minimum detected value, and variance-filtered before integration.
